# Nucleocytoplasmic transport rates are regulated by cellular processes that modulate GTP availability

**DOI:** 10.1101/2023.12.29.573651

**Authors:** Kelsey L. Scott, Charles T. Halfmann, Allison D. Hoefakker, Purboja Purkayastha, Ting Ching Wang, Tanmay P. Lele, Kyle J. Roux

## Abstract

Nucleocytoplasmic transport (NCT), the facilitated diffusion of cargo molecules between the nucleus and cytoplasm through nuclear pore complexes (NPCs), enables numerous fundamental eukaryotic cellular processes. Ran GTPase uses cellular energy in the direct form of GTP to create a gradient across the nuclear envelope (NE) that drives the majority of NCT. We report here that changes in GTP availability resulting from altered cellular physiology modulate the rate of NCT, as monitored using synthetic and natural cargo, and the dynamics of Ran itself. Cell migration, cell spreading and/or modulation of the cytoskeleton or its connection to the nucleus alter GTP availability and thus rates of NCT, regulating RNA export and protein synthesis. These findings support a model in which changes in cellular physiology that alter GTP availability can regulate the rate of NCT, impacting fundamental cellular processes that extensively utilize NCT.

**Summary:** Changes in the availability of cellular GTP resulting from physiologically relevant processes, including cell migration and cell spreading, alter the rates of Ran-dependent nuclear import and export. Altered rates of nucleocytoplasmic transport regulate RNA localization and protein synthesis.

## Introduction

Separation of cytoplasmic and nucleoplasmic compartments, a distinguishing feature for all eukaryotes, is critical to fundamental cellular processes including genome protection, regulation of transcription and translation, and myriad signaling events. The nuclear envelope (NE), a specialized extension of the endoplasmic reticulum, surrounds the genome with a double-membraned physical barrier compartmentalizing the nucleus from the cytoplasm. NCT occurs through NPCs embedded within the NE. NPCs are well-conserved structures capable of permitting regulated transit of cargo as large as ∼39 nm (Pante and Kann, 2002) and excluding unregulated transit of molecules larger than ∼9 nm (Gorlich and Kutay, 1999). Regulated transport through the NPCs depends on importins and exportins (Ding and Sepehrimanesh, 2021). These transport receptors selectively bind to specific cargo and shuttle them through the hydrophobic FG-repeat barrier; the transport culminates in dissociation and release of cargo into the other compartment (Aramburu and Lemke, 2017; Tan et al., 2018). For most cargo regulated by NCT, the binding and/or dissociation of transport receptors to cargo is catalyzed by the small GTPase Ran. Nuclear Ran is primarily bound to GTP while cytoplasmic Ran is primarily bound to GDP (Mattaj and Englmeier, 1998). There is a steep gradient of nuclear RanGTP compared to the cytoplasm that is required for Ran-dependent NCT (Izaurralde et al., 1997).

The transport of cargo across the NE through NPCs does not directly require energy; however, the maintenance of the Ran gradient is an energy-dependent process (Englmeier et al., 1999; Lyman et al., 2002; Ribbeck et al., 1999; Schwoebel et al., 2002). RanGTP exits the nucleus with exportins and bound cargo as well as with importins as they recycle back to the cytoplasm. Once in the cytoplasm, the Ran converts the bound GTP to GDP in a process facilitated by RanGAP and Ran-binding proteins (Bischoff et al., 1994; Bischoff et al., 1995; Mahajan et al., 1997; Matunis et al., 1996; Yokoyama et al., 1995). This net flux of Ran out of the nucleus and conversion to RanGDP that occurs as a result of NCT would rapidly deplete the Ran gradient and grind the system to a halt. However, RanGDP is shuttled back into the nucleus by NTF2 (Ribbeck et al., 1998) where it binds the Ran GEF, RCC1, leading to release of the bound GDP and enabling exchange for the more abundant GTP (Klebe et al., 1995b). The binding affinity of Ran for GTP is lower than for GDP (Klebe et al., 1995b), but the ratio of nuclear GTP:GDP is typically favorable for nuclear Ran to be predominantly GTP bound (Kalita et al., 2021). This last step is the primary energy-dependent step in NCT with GTP being the immediate energy source.

It has been estimated that approximately 10^5^ NCT events occur per second in a typical mammalian cell (Gorlich et al., 2003; Smith et al., 2002) with each event requiring consumption of at least one GTP to recharge the system and maintain the Ran gradient. Many critical cellular processes are GTP dependent including protein synthesis, microtubule dynamics, DNA and RNA synthesis, vesicle transport, cytoskeletal regulation, and G-protein signaling. Previous studies have explored how variations in the availability and even compartmentalization of GTP can regulate the activity of small GTPases like Rac or Rho, impacting actin dynamics and regulating cellular processes such as the formation of membrane protrusions involved in cell migration (Bianchi-Smiraglia et al., 2021; Wawrzyniak et al., 2013). Similarly, the GTPase dynamin superfamily has been shown to be sensitive to changes in GTP availability, altering cellular membrane dynamics at the cell surface and mitochondria (Boissan et al., 2014). Replenishment of GTP by adding a phosphate to GDP is predominantly enabled by nucleoside diphosphate kinase-mediated phosphate shuttling from ATP (Georgescauld et al., 2020), thus the pools of ATP and GTP are intimately connected. Processes that typically consume substantial amounts of available cellular energy include protein synthesis (Lindqvist et al., 2018), generation of ionic gradients (Harris et al., 2012), as well as maintenance of the cytoskeleton and its use in force generation (DeWane et al., 2021).

Although the energy dependence of NCT is well established, there is little information available as to how physiologically relevant variations in cellular energy, and more specifically availability of GTP, impact rates of NCT. Here we have employed a live-cell GTP sensor and multiple reporters of NCT, including reporters of the dynamics of Ran itself, to investigate how the rates of NCT are altered by both artificially induced fluctuations in available GTP and physiologically relevant processes that consume cellular energy. Furthermore, we begin to explore the biological impacts of altered NCT rates, observing changes in RNA export and protein synthesis, processes which extensively utilize Ran GTPase-dependent NCT.

## Results

### Modulating GTP availability alters rates of NCT in living cells

To assess how GTP availability impacts NCT, we initially validated our chosen tools to measure GTP levels and NCT rates in live cells. We utilized hTERT-immortalized human diploid fibroblasts (BJ-5ta) throughout these studies unless otherwise indicated. To assess NCT, we employed the live-cell light inducible nuclear localization signal (LINuS) reporter that enables measurement of nuclear import and export rates (Niopek et al., 2014). LINuS is based on a fusion between mCherry and a strong Importin α1/β1 nuclear localization sequence (NLS) that is caged in an inactive conformation via a light sensitive LOV2 domain and a weaker yet constitutively active CRM1-mediated nuclear export sequence (NES), leading to cytoplasmic localization of the reporter (Fig. 1 A). Exposure to 488 nm light reversibly uncages the NLS, enabling the NCT-mediated translocation of the LINuS reporter into the nucleus. The NLS is re-caged upon removal of 488 nm light exposure, enabling the constitutively active NES to mediate NCT-based export of the LINuS reporter out of the nucleus (Fig. 1 B). This inducible accumulation and loss of nuclear LINuS can be used to measure active import and export, respectively (Fig. 1 C). We assessed if LINuS negatively impacts NCT due to an increase in cargo competing for importins by overexpressing a GFP fused to three tandem NLS (GFP-NLSx3) that uses the same importins as LINuS and observed no significant change in import or export rates for LINuS, suggesting that expression of LINuS does not appreciably alter NCT (Fig. S1 A). To directly monitor cellular levels of available GTP, we used the live-cell GTP reporter system, GEVAL, that is based on a circularly permutated YFP fused to variable GTP-binding domains (Bianchi-Smiraglia et al., 2017). Binding to GTP changes the fluorescent properties of the YFP, altering the fluorescence intensity when excited at 488 nm but not 405 nm, allowing for ratiometric imaging for evaluating intracellular GTP levels independent of GEVAL expression level. To measure available cellular GTP, we utilized GEVAL30, a GEVAL variant that can bind GTP at physiologically relevant concentrations. To ensure changes in GEVAL fluorescence are GTP-dependent, we also utilized GEVALNull, a control reporter which cannot bind GTP. An additional advantage to the GEVAL system is the ability to measure subcellular concentrations of GTP. Since the nuclear pool of GTP is utilized to maintain the Ran gradient, we measured GEVAL intensity in the nucleus. To validate these tools, we reduced intracellular GTP levels with mycophenolic acid (MPA; Fig. 1, D and E) that blocks GTP precursor synthesis but retains the purine salvage pathway, thus it does not fully deplete cellular GTP (Ransom, 1995). GTP depletion by MPA resulted in a considerable decrease in NCT as measured with LINuS, which could be rescued by guanosine supplementation (Fig. 1 F). We next tested if increased availability of cellular GTP would alter NCT by inhibiting protein synthesis, a cellular process that consumes considerable GTP (Lindqvist et al., 2018). We observed an increase in GTP levels (Fig. G and H) and rates of NCT (Fig. 1 I) 2hr after inhibition of protein synthesis with cycloheximide. We also validated that ATP depletion profoundly reduced rates of NCT (Fig. 1 J), as previously reported (Schwoebel et al., 2002). Collectively, these experiments align with prior studies on the bioenergetic dependence of NCT, validate our use of GEVAL and LINuS to monitor available cellular GTP and rates of NCT, and that altering a cellular process that directly consumes GTP alters rates of NCT.

**Figure 1:**
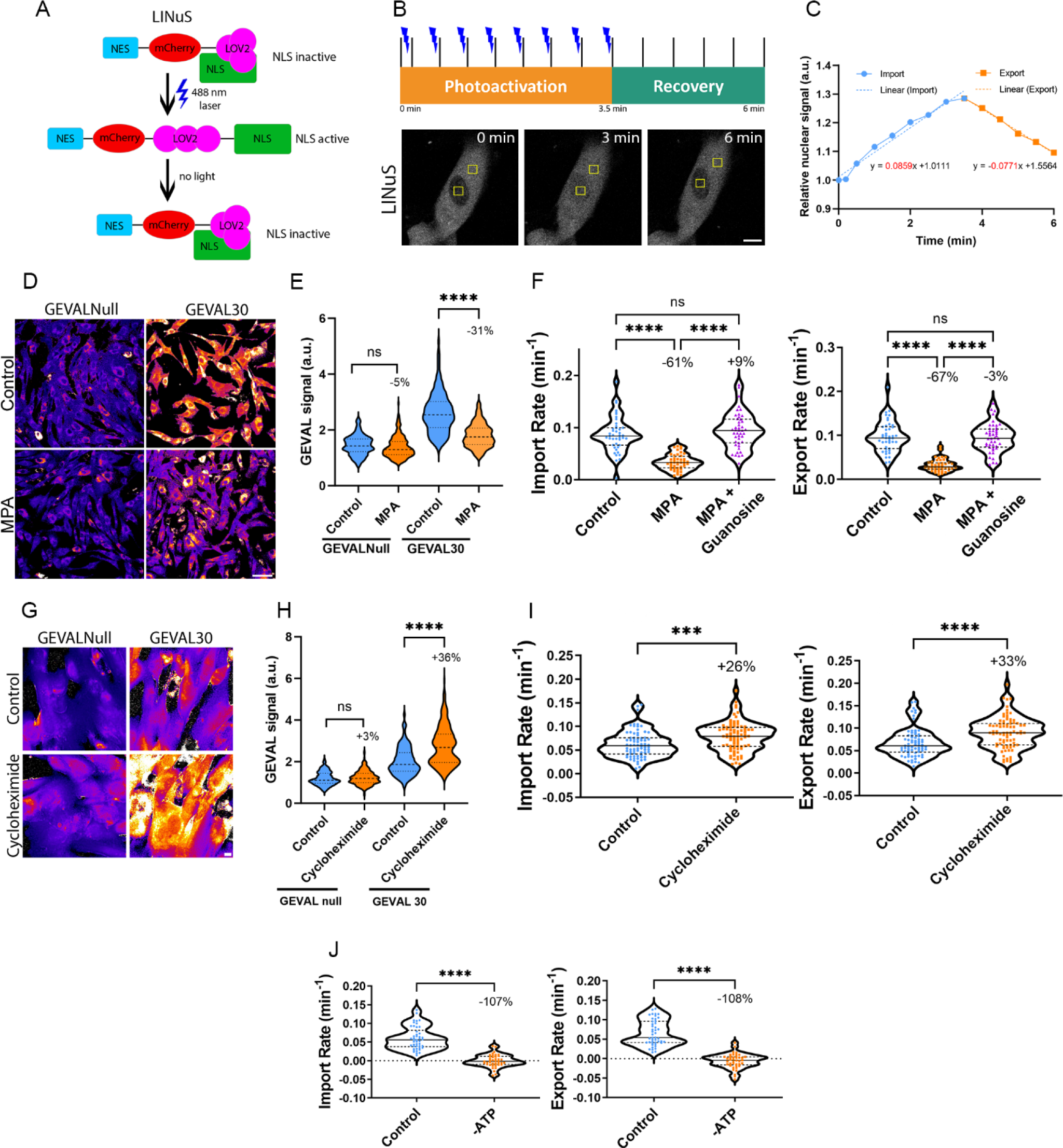
Nucleocytoplasmic transport rates are sensitive to levels of GTP. **(A)** Model of photoactivatable NCT reporter, LINuS. **(B)** Timeline of LINuS photoactivation and recovery, with representative IF images in BJ-5ta cells stably expressing LINuS during photoactivation and recovery. Yellow boxes denote areas of measurement. **(C)** Representative graph of LINuS nuclear localization with trend lines, the slopes of which provide import or export rates. **(D)** Ratiometric images (405nm/488nm) of BJ-5ta cells expressing either GEVALNull or GEVAL30 treated with MPA. **(E)** Quantification of the GEVAL ratiometric signal (405nm/488nm) for each cell expressing GEVALNull (control n=186, MPA n=223) or GEVAL30 (control n=175, MPA n=264). Results are from two independent replicates. **(F)** Import and export rates of BJ-5ta cells expressing LINuS treated with MPA to deplete GTP (control n=43, MPA n=43), then rescued with guanosine supplementation (MPA + guanosine n=44). Results are from two independent replicates. **(G)** Ratiometric images (405nm/488nm) of BJ-5ta cells expressing either GEVALNull or GEVAL30 treated with cycloheximide. **(H)** Quantification of cells from G (GEVALNull control n=203, cycloheximide n=232, GEVAL30 control n=197, cycloheximide n=198; 3 independent experiments). **(I)** Import and export rates of BJ-5ta cells expressing LINuS treated with cycloheximide (control n=74, cycloheximide n=82; 2 independent replicates). **(J)** Quantification of GEVAL ratiometric signal (405nm/488mn) in BJ-5ta cells expressing GEVALNull (control n=73, GTP n=77) or GEVAL30 (control n=138, GTP n=122) that were treated with GTP. Results are from two independent replicates. **(K)** Import and export rates of BJ-5ta cells expressing LINuS treated with GTP (control n=99, GTP n=131; 3 independent replicates). **(L)** Import and export rates of BJ-5ta cells depleted of ATP with sodium azide and 2-deoxyglucose (control n=43, - ATP n=43; two independent replicates). Significance calculated using unpaired t-test (E, H, I, J), one-way ANOVA with Tukey’s post hoc (F). ns P>0.05, P***<0.001, P****<0.0001. Scale bars 10 µm.

### Cell spreading and migration reduce available GTP and rates of NCT

Inhibiting protein synthesis increases available GTP and NCT rates, but this inhibition is unlikely to occur naturally. Next, we assessed if naturally occurring changes in cellular physiology would predictably alter available GTP and thus rates of NCT. A significant portion of cellular energy consumption is for the maintenance and use of the cytoskeleton to generate cellular forces (DeWane et al., 2021). Cells grown on stiffer substrates are typically more spread and generate more cellular tension than do those on soft substrates (Solon et al., 2007; Yeung et al., 2005). We grew cells on fibronectin-coated polyacrylamide hydrogels with substrate rigidities of 1, 22, 46, or 308 kPa (Fig. 2 A). The nuclear volumes were similar for cells on all tested substrate rigidities (Fig. 2 B), but cells on 1 kPa substrates were markedly less spread with significantly increased nuclear height compared to those on the stiffer substrates, consistent with previous reports (Fig. 2 C) (Lovett et al., 2013). Cells became increasingly spread as substrate rigidity increased, with the spreading plateauing at 46 kPa (Fig. 2 D). Compared to cells grown on 46 or 308 kPa substrates, cells on both 1 and 22 kPa substrates exhibited significantly elevated levels of GTP (Fig. 2 E) and rates of NCT (Fig. 2 F). To further explore how cell spreading regulates NCT, we rounded the cells by trypsinization, followed by re-plating to initiate cell spreading. Two hr post-plating, cells were attached but rounded compared to 24-hr post plating when cells were spread (Fig. 2, G and H). We observed a profound decrease in NCT in the spread cells at 24 hr compared to the rounded cells 2 hr post-plating (Fig. 2 H). To assess a more physiologically relevant change in cell spreading, monolayers of confluent MCF10A epithelial cells were ‘scratch wounded’ to induce cell migration towards the cell-denuded area (Fig. 2 J) (Tse et al., 2012). Available GTP levels (Fig. 2 K) and NCT rates (Fig. 2 L) were measured in cells 2 hr post-scratch as they began to migrate into the denuded area (scratch) and compared to cells > 600 microns distal from the scratch (middle). Both available GTP and NCT rates were significantly decreased in cells that had flattened out and were migrating into the denuded area, compared to those within the monolayer that were less spread or motile. Collectively, these results demonstrate that naturally-occurring changes in cell behavior, including spreading induced by substrate rigidity or migration, can alter levels of available GTP and rates of NCT.

**Figure 2:**
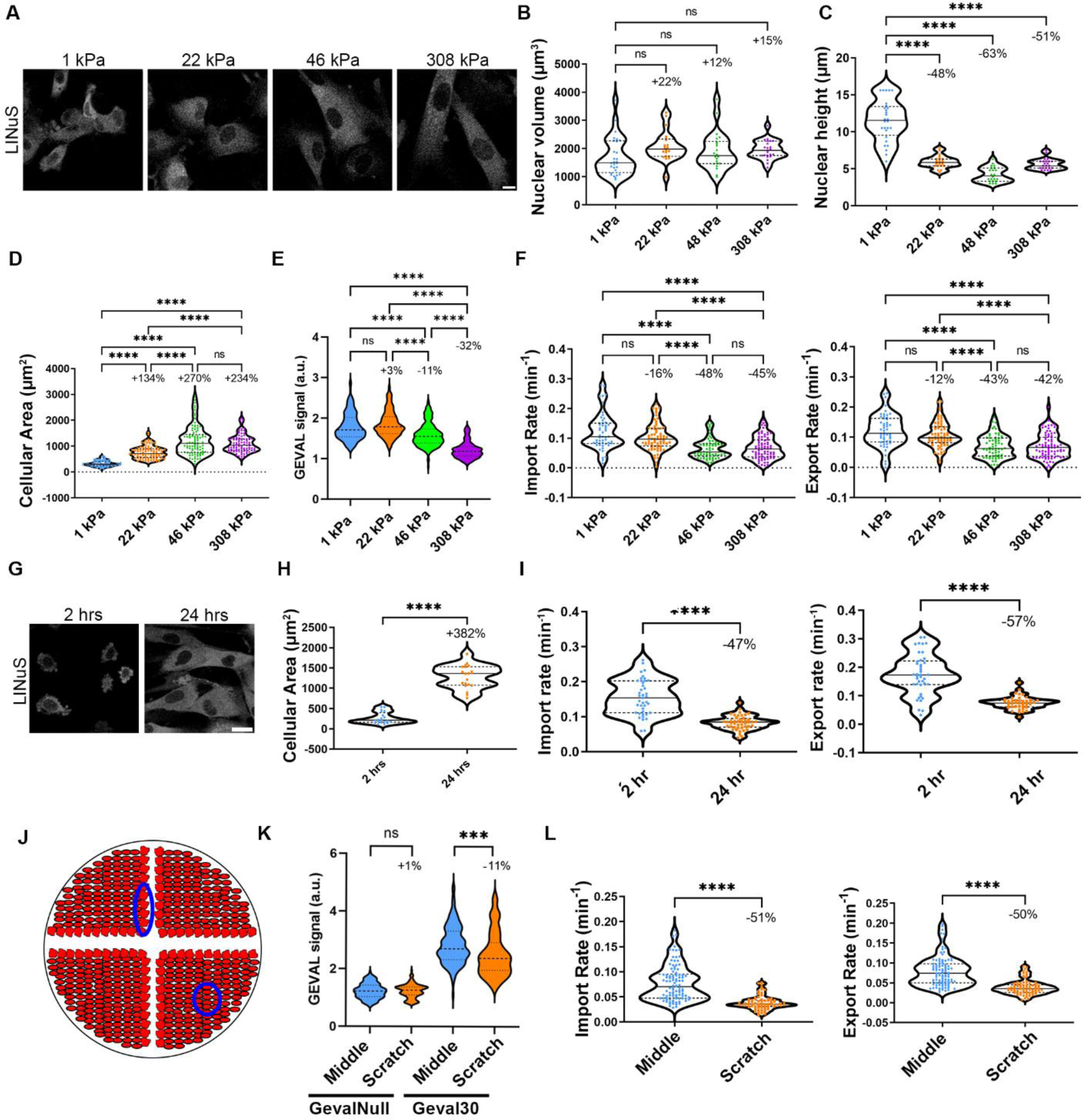
Cell spreading regulates levels of GTP and rates of NCT. **(A)** Representative confocal images of BJ-5ta cells expressing LINuS plated on polyacrylamide hydrogels with rigidities of 1, 22, 46 or 308 kPa. **(B)** Nuclear volume (1kPa n=23, 22 kPa n=22, 46 kPa n=21, 308 kPa n=28; two independent replicates), **(C)** nuclear height (1kPa n=23, 22 kPa n=22, 46 kPa n=21, 308 kPa n=28; two independent replicates), or **(D)** cellular area (1kPa n=81, 22 kPa n=93, 46 kPa n=93, 308 kPa n=93; two independent replicates) of BJ-5ta cells plated on polyacrylamide hydrogels of 1, 22, 46, or 308 kPa rigidities. **(E)** Quantification of GEVAL30 signal (405nm/488nm) in BJ-5ta cells expressing GEVAL30 in cells plated on polyacrylamide hydrogels at stiffnesses of 1 (n=152), 22 (n=178), 46 (n=197), or 308 kPa (n=172). Results are from two independent replicates. **(F)** Import and export rates of BJ-5ta cells expressing LINuS plated on polyacrylamide hydrogels with rigidities of 1 (n=52), 22 (n=72), 46 (n=64), or 308 kPa (n=80) across four independent replicates. **(G)** Representative confocal images of BJ-5ta cells expressing LINuS 2 or 24 hr after being plated. **(H)** Cellular area of BJ-5ta cells that had been plated for 2 (n=20) or 24 hr (n=20) across two independent replicates. **(I**) Import and export rates of BJ-5ta cells expressing LINuS either 2 or 24 hr after being plated (2 hr n=41, 24 hr n=44; two independent replicates). **(J)** Cartoon of MCF10A cells grown into a monolayer, then scratch wounded. Cells were either imaged along the scratch wound (scratch) as they migrated into the denuded space or ∼600 microns into the monolayer adjacent from the scratch edge (middle). Areas of interest depicted by blue ovals. **(K)** Quantification of GEVAL signal (405nm/488nm) in MCF10A cells expressing GEVALNull or GEVAL30 within the monolayer (middle, GEVALNull n=147, GEVAL30 n=138) or edge of scratch (scratch, GEVALNull n=153, GEVAL30 n=106) two hr after scratch. Results are from two independent replicates. **(L)** Import and export rates of MCF10A cells expressing LINuS within the monolayer (middle, n=98) or edge of scratch (scratch, n=72) two hr after scratch, across three independent replicates. Significance calculated using unpaired t-test (H, I, K, L), one-way ANOVA with Tukey’s post hoc (D, E, F), or one way-ANOVA with Dunnett’s post hoc (B, C). ns P>0.05, P***<0.001, P****<0.0001. Scale bars 10 µm.

### Actin and microtubule networks and their connection to the NE mediate GTP consuming processes that regulate NCT rates

Changes in cell shape and migration require energy consuming force-generating structures such as the actin or microtubule networks. Therefore, we explored how perturbation of those structures in spread cells impact GTP availability and rates of NCT. Depolymerization of actin networks in BJ-5ta cells (Fig. S2 A) with cytochalasin B rapidly increased GTP availability and rates of NCT within 15 min (Fig. 3, A, B and C). A similar effect on NCT was observed with actin depolymerizing latrunculin B within 10 min (Fig. 3 C). Depolymerization of the microtubule networks with nocodazole (Fig. S2 B) resulted in a similar increase in rates of NCT (Fig. 3 C). Depolymerization of the actin cytoskeleton in MCF10A cells with cytochalasin B similarly increased available GTP and NCT rates (Fig. 3, D and E). In contrast, perturbation of the vimentin intermediate filament network by siRNA (siVimentin; Fig. S2 C) failed to alter either GTP levels or rates of NCT (Fig. 3, F and G).

**Figure 3:**
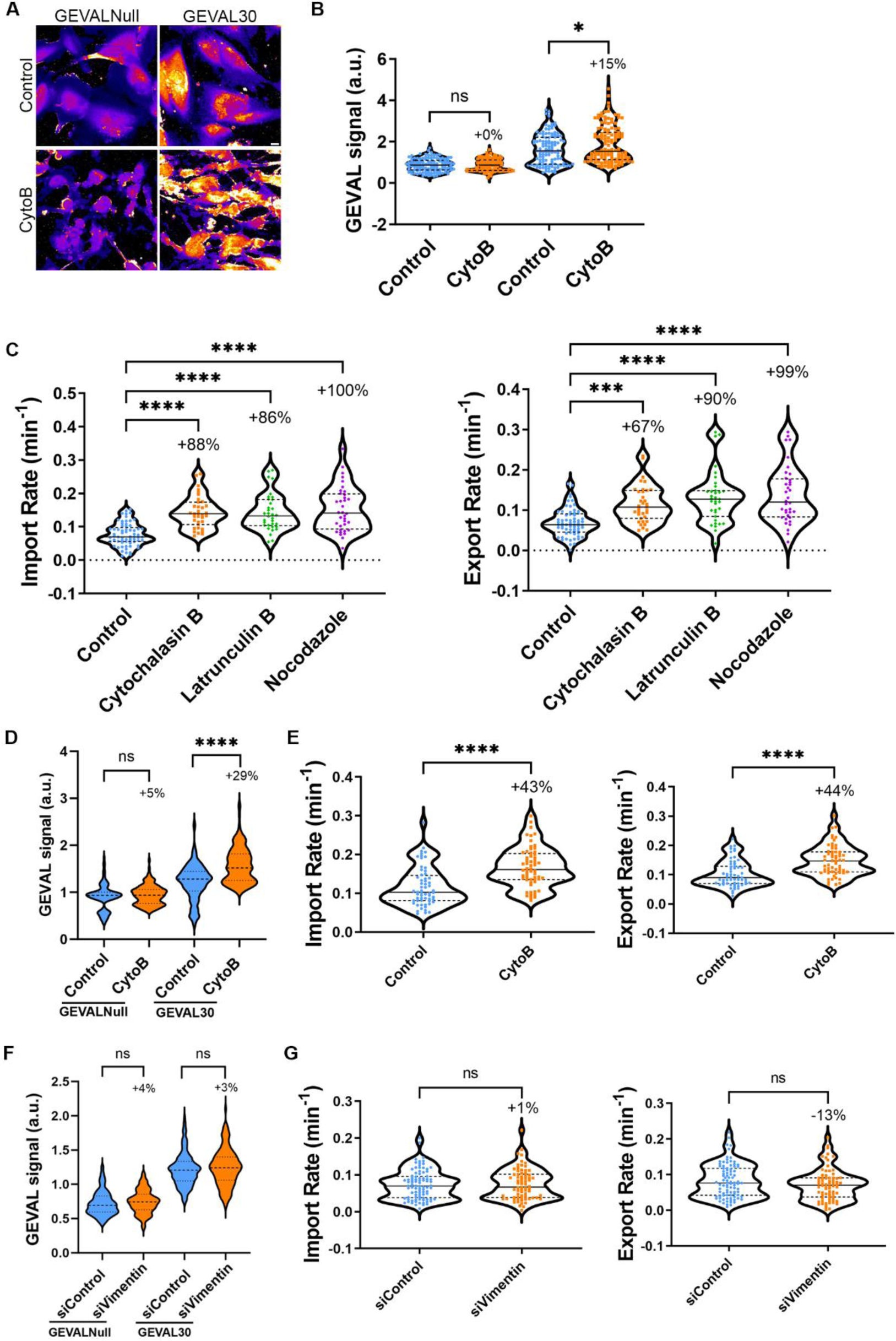
Disruption of actin and microtubule cytoskeleton enhances GTP availability and rates of NCT. **(A)** Representative ratiometric images (405nm/488nm) of BJ-5ta cells expressing either GEVALNull or GEVAL30 treated with cytochalasin B. **(B)** Quantification of GEVAL ratiometric signal (405nm/488nm) in BJ-5ta cells expressing GEVALNull or GEVAL30 treated with cytochalasin B (GEVALNull control and cytochalasin B, n=121 and n=135 respectively; GEVAL30 control and cytochalasin B, n=117 and n=130, respectively. Results are from two independent replicates. **(C)** Import and export rates of BJ-5ta cells expressing LINuS treated with cytochalasin B, latrunculin B, or nocodazole (control n= 65, cytochalasin B n=35, latrunculin B n=34, nocodazole n=36). Results are from three independent replicates. **(D)** Quantification of GEVAL ratiometric signal (405nm/488nm) in MCF10A cells expressing GEVALNull or GEVAL30 treated with cytochalasin B (GEVALNull control and cytochalasin B, n=101 and n=123 respectively; GEVAL30 control and cytochalasin B, n=134 and n=157, respectively). Results are from three independent replicates. **(E)** Import and export rates of MCF10A cells expressing LINuS treated with cytochalasin B (control n=60, cytochalasin B n=60; results from three independent replicates). **(F)** Quantification of GEVAL ratiometric signal (405nm/488nm) in BJ-5ta cells expressing GEVALNull or GEVAL30 depleted of vimentin (GEVALNull siControl and siVimentin, n=128 and n=136, respectively; GEVAL30 siControl and siVimentin, n=179 and n=118, respectively). Results are from three independent replicates. **(G)** Import and export rates of BJ-5ta cells expressing LINuS depleted of vimentin (siControl n=89, siVimentin n=74; 3 independent replicates). Significance calculated using unpaired t-test (B, D-G), or one-way ANOVA with Dunnett’s post hoc (C). ns P>0.05, P***<0.001, P****<0.0001. Scale bar 10 μm.

We next assessed if the connections between the cytoskeleton and the nucleus participate in the cellular tension-based regulation of NCT. The LINC complex, a primary mediator of these associations (Hoffman et al., 2020; Jahed et al., 2016; Kuhn and Capelson, 2019) can be perturbed by co-depletion of Suns1 and 2 (Figure S2 D) which leads to an increase in GTP levels (Fig. 4 A) and rates of NCT (Fig. 4 B). Individual depletion of Sun1 (Figure S2 E) failed to significantly increase these levels for either, and if anything resulted in a slight reduction in NCT (Fig. 4, A and B), whereas depletion of Sun2 (Figure S2 F) recapitulated the results of simultaneous co-depletion (Fig. 4, A and B). This suggests that in BJ-5ta cells, Sun2 is the primary constituent of LINC complex that mediates these effects on GTP availability and NCT rates. The impact of LINC complex perturbation on NCT rates was investigated via expression of a dominant negative lumenal fragment of Sun1 (SS-GFP-Sun1L-KDEL; Fig. S2 G) that outcompetes endogenous Suns for binding to the KASH-domain nesprin proteins (Crisp et al., 2006). Compared to cells expressing a control (SS-GFP-KDEL), the LINC perturbed cells exhibited increased rates of NCT (Fig. 4 C). We next assessed the roles of individual nesprins in mediating these LINC complex effects on NCT. Nesprins1 and 2 can be extremely large proteins with a multitude of splice isoforms, but function in part to directly tether the actin cytoskeleton and indirectly tether the microtubules by binding to kinesin and dynein motor proteins to the nucleus (Gundersen and Worman, 2013; Luxton et al., 2010; Zhang et al., 2001). Nesprin3 is a smaller KASH domain protein that binds to the cytolinker plectin and is primarily associated with cytoplasmic intermediate filament association at the nuclear surface (Wilhelmsen et al., 2005). Depletion of either Nesprin1 or 2 using siRNAs (siNesprin1 or siNesprin2, respectively), but not Nesprin3 (siNesprin3; Fig. S2, H, I and J), increased levels of available GTP (Fig. 4 D) and increased rates of NCT (Fig. 4 E). To control for possible siRNA off-target effects, we performed LINuS experiments using CRISPR interference (KRAB-dCas9-IRES-LINuS) to deplete these proteins (Fig. S2 K). Thus far, we have observed that cell spreading and migration, supported by actin and microtubule networks that are connected to the nucleus via the LINC complex, regulate the levels of available GTP and modulate rates of NCT. These findings are logical and perhaps should be expected from the perspective of bioenergetics; however, since cell spreading and/or migration can increase forces on the nucleus, they appear to contradict a recent study reporting that NCT is enhanced by increased forces on the nucleus that mechanically dilate NPCs (Andreu et al., 2022). In support of our findings, it has been reported that the translocation capacity of NPCs considerably exceeds the actual rates of NCT and hypothesized that transportin-cargo binding and/or release, and not the size or permeability of the NPCs, is the primary rate limiting step in NCT (Ribbeck and Gorlich, 2001). To ensure that our discrepant findings are not an artifact of the LINuS reporter, we utilized methods similar to those in the prior study (Andreu et al., 2022), namely transient transfection of LINC perturbing GFP-KASH in mouse embryonic fibroblasts (MEFs) expressing an alternate photoactivatable NCT reporter called LEXY (Niopek et al., 2016). We observed that LINC perturbation resulted in enhanced NCT of LEXY, similar to our findings with LINuS (Fig. S3, A, B and C). Collectively, these results support a model for adherent spread cells in which cell-substrate interactions and the actin and microtubule networks and their connection to the nucleus enable the generation of cellular forces that consume energy such that levels of GTP, and thus, rates of NCT are reduced.

**Figure 4:**
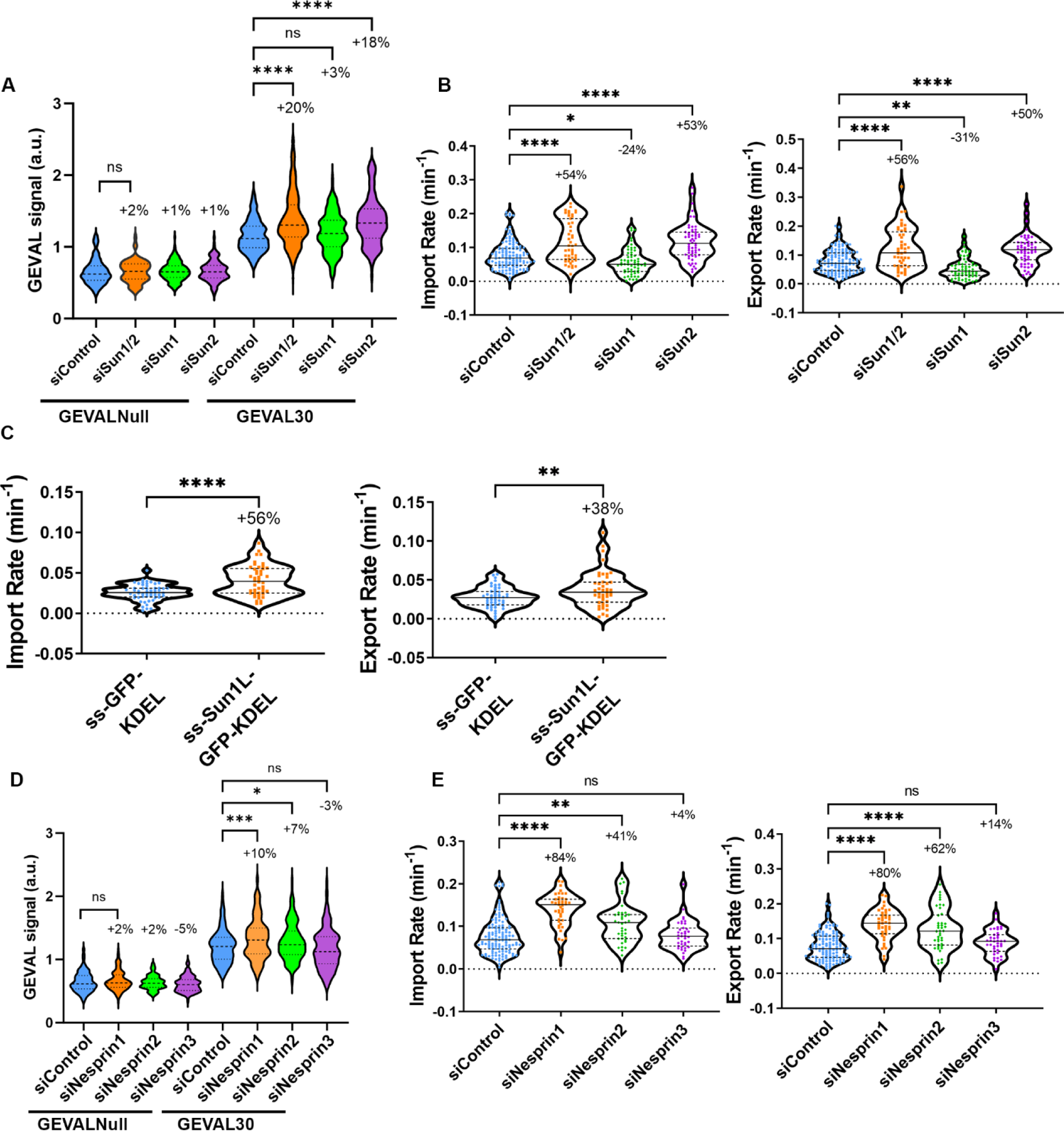
Disruption of the LINC complex enhances GTP availability and rates of NCT. **(A)** Quantification of GEVAL ratiometric signal (405nm/488nm) in BJ-5ta cells expressing GEVALnull or GEVAL30 and co-depleted of Sun1 and Sun2 (Sun1/2), Sun1, or Sun2 (GEVALNull siControl n=130, siSun1/2 n=115, siSun1=173, siSun2=128. GEVAL30 siControl n=173, siSun1/2 n=203, siSun1 n=227, siSun2 n=153). Results are from three independent replicates. **(B)** Import and export rates of BJ-5ta cells stably expressing LINuS and co-depleted of Sun1 and Sun2 (Sun1/2), Sun1, or Sun2. siControl n=125, siSun1/2 n=47, siSun1 n=61, siSun2 n=58. Results are from three independent replicates. **(C)** Import and export rates of MCF10A cells stably expressing LINuS and transiently transfected with ss-GFP-KDEL (n=53) or ss-Sun1L-GFP-KDEL (n=46) from three independent replicates. **(D)** Quantification of GEVAL ratiometric signal (405nm/488nm) in BJ-5ta cells expressing GEVALnull or GEVAL30 depleted of Nesprin1, Nesprin2, or Nesprin3 (GEVALNull siControl n=138, siNesprin1 n=144, siNesprin2 n=107, siNesprin3 n=132; GEVAL30 siControl n=180, siNesprin1 n=154, siNesprin2 n=190, siNesprin3 n=144). Results are from three independent replicates. **(E)** Import and export rates of BJ-5ta cells stably expressing LINuS and depleted of Nesprin1, Nesprin2, or Nesprin3 (siControl n=125, siNesprin1 n=43, siNesprin2 n=39, siNesprin3 n=39; 3 independent replicates). Significance calculated using t-test (C), or one-way ANOVA with Dunnett’s post hoc (A-B, D-E). ns P>0.05, P*<0.05, P**<0.01, P***<0.001, P****<0.0001.

### Conditions that alter GTP availability similarly change rates of induced glucocorticoid receptor import

To expand our findings beyond the more artificial reporters of LINuS or LEXY that rely on minimal NLS and NES sequences specific to importin α/β and Crm1-mediated NCT and to assess if the impacts of altered GTP on NCT rates are specific to transport of those reporters, or instead more globally impact Ran-mediated transport, we utilized inducible import of glucocorticoid receptor (GR). GR is a transcription factor whose nuclear import is mediated by Importin7, in contrast to Importin α/β-based for LINuS, and is inducible with dexamethasone (Hakim et al., 2013). GR-GFP was stably expressed in BJ-5ta cells and nuclear import was induced by exposure to dexamethasone. GR-GFP utilization of an importin distinct from LINuS was confirmed by inhibiting LINuS import with importazole, an inhibitor of importin-β (Fig. 5, A and B), which had no impact on the induced import of GR-GFP (Fig. 5, C and D). Utilizing some of the same modulators of GTP availability as was done previously with LINuS, we observed a significant increase in GR import when cells were rounded at 2 hr compared to spread at 24 hr (Fig. 5 E) or when cells were depleted of Sun2 (Fig. 5 F). These findings support our hypothesis that the impact of GTP availability on NCT is not limited to specific transportins, but predicted to affect all Ran gradient-dependent NCT.

**Figure 5:**
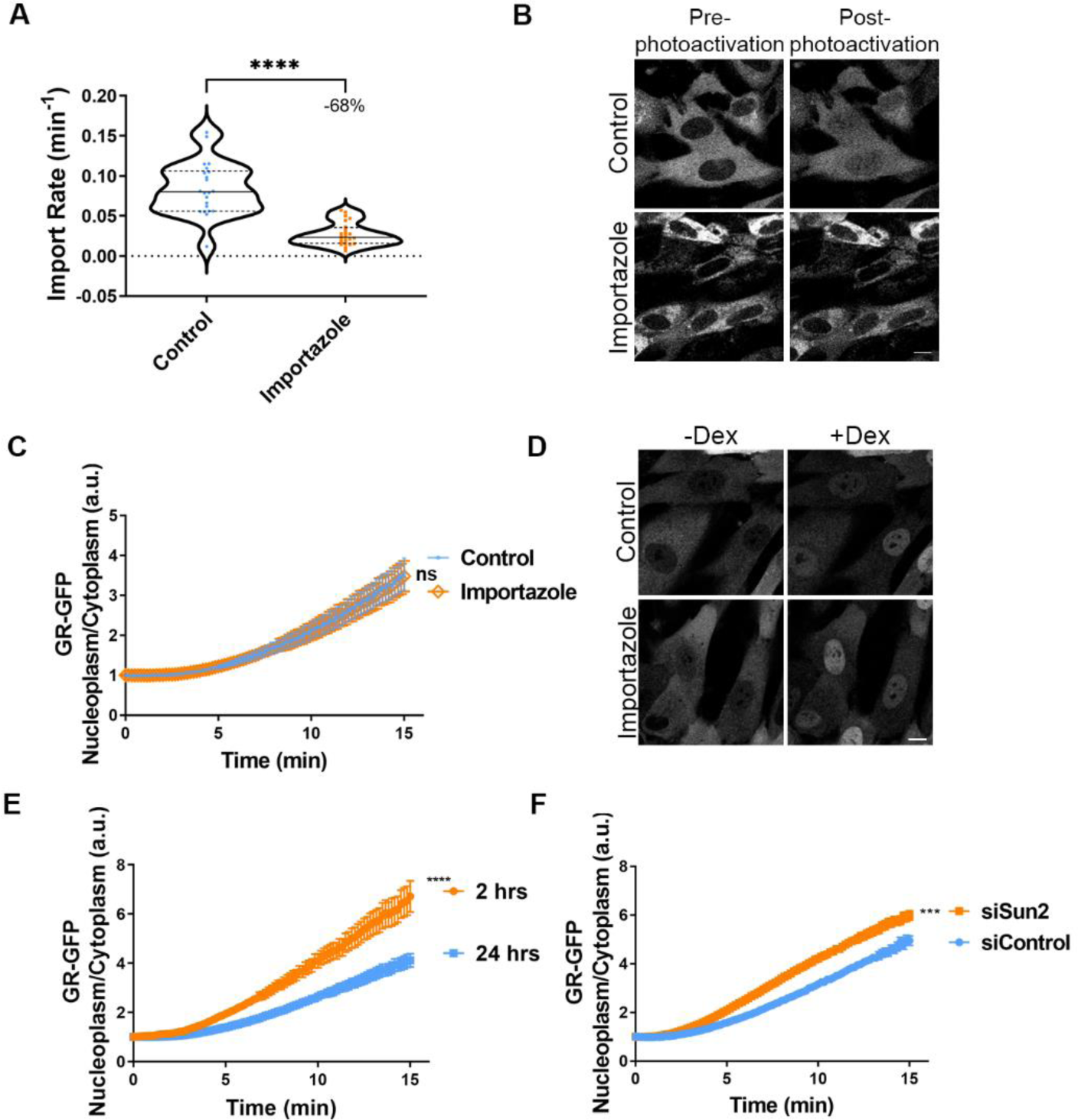
Conditions that alter GTP availability similarly modulate rates of induced GR import. **(A)** Import rates of BJ-5ta cells expressing LINuS and treated with importazole (control n=22, importazole n=24; two independent replicates). **(B)** Representative confocal images of pre-photoactivation and 3.5 min post-photoactivation of LINuS in BJ-5ta cells treated with importazole. **C)** Quantitative measurements of dexamethasone-induced import of GR-GFP in BJ-5ta cells with or without importazole (Control n=22; importazole n=25; 2 independent replicates), with representative confocal images **(D)** of cells with either no dexamethasone treatment (-Dex) or 15 min after dexamethasone treatment (+Dex). **(E-F)** Graphs of dexamethasone-induced nuclear import of GR-GFP in BJ-5ta cells **(E)** plated for either 2 (n=36) or 24 hr (n=38; 2 independent replicates) or **(F)** depleted of Sun2 (siControl n=105, siSun2 n=107; 2 independent replicates). Significance calculated using unpaired t-test (A, C, E, F). P***<0.001, P****<0.0001. Scale bars 10 µm. Error bars represent ±SEM.

### Ran GTPase dynamics and localization are modulated by levels of available GTP

To investigate if, in addition to cargo and transportins, Ran GTPase itself exhibits altered rates of nuclear entry and exit based on changes in GTP availability, we examined the dynamics of Ran. We utilized fluorescence loss in photobleaching (FLIP) of cytoplasmic GFP-Ran to measure the subsequent loss of fluorescence from the nucleus, where it predominantly accumulates under normal conditions. We observed a significant increase in the loss of GFP-Ran from the nucleus with conditions that increase GTP availability and rates of NCT, such as Sun2 depletion (Fig. 6, A, B and C), actin depolymerization (Fig. 6, D and E), cell rounding (Fig. 6, F and G), or inhibition of protein synthesis (Fig. 6, H and I). During the course of the GFP-Ran mobility studies, we observed that the enhanced rate of nuclear exit of Ran driven by elevated levels of GTP coincided with an increase in the steady-state localization of Ran in the cytoplasm. Indeed, we observed that under a prolonged period of enhanced GTP availability and NCT, such as occurs with the depletion of Sun2, total levels of endogenous Ran do not change but Ran is considerably more cytoplasmic (Fig. 7, A, B and C). Using a more rapid approach to enhance GTP levels by the addition of cytochalasin B, we can also observe an abrupt shift in the localization of endogenous Ran (Fig. 7 D) as well as an exogenous GFP-Ran towards the cytoplasm, both in BJ-5ta (Fig. 7, E, F and G, Movie S1 and S2) and MCF10A cells (Fig. 7, H, I and J) compared to untreated cells. We also observed a rapid shift from cytoplasmic to nuclear localization of GFP-Ran within about 2 hr after cell plating that correlated with an increase in cellular spreading area (Fig. 7, K and L). To test if Ran accumulates in the cytoplasm under conditions of enhanced NCT because nuclear reimport by NTF2 is a rate limiting factor in its nucleocytoplasmic exchange, as has been previously proposed (Ribbeck et al., 1998; Steggerda et al., 2000), we stably overexpressed mCherry-NTF2 and observed a partial rescue of Ran localization following perturbation of the actin cytoskeleton (Fig. 7, M and N) or disruption of the LINC complex (Fig. 7 O). Together, these data suggest that the rates of Ran shuttling between the nucleus and cytoplasm are increased under conditions of enhanced NCT that occurs coincident with elevated GTP.

**Figure 6:**
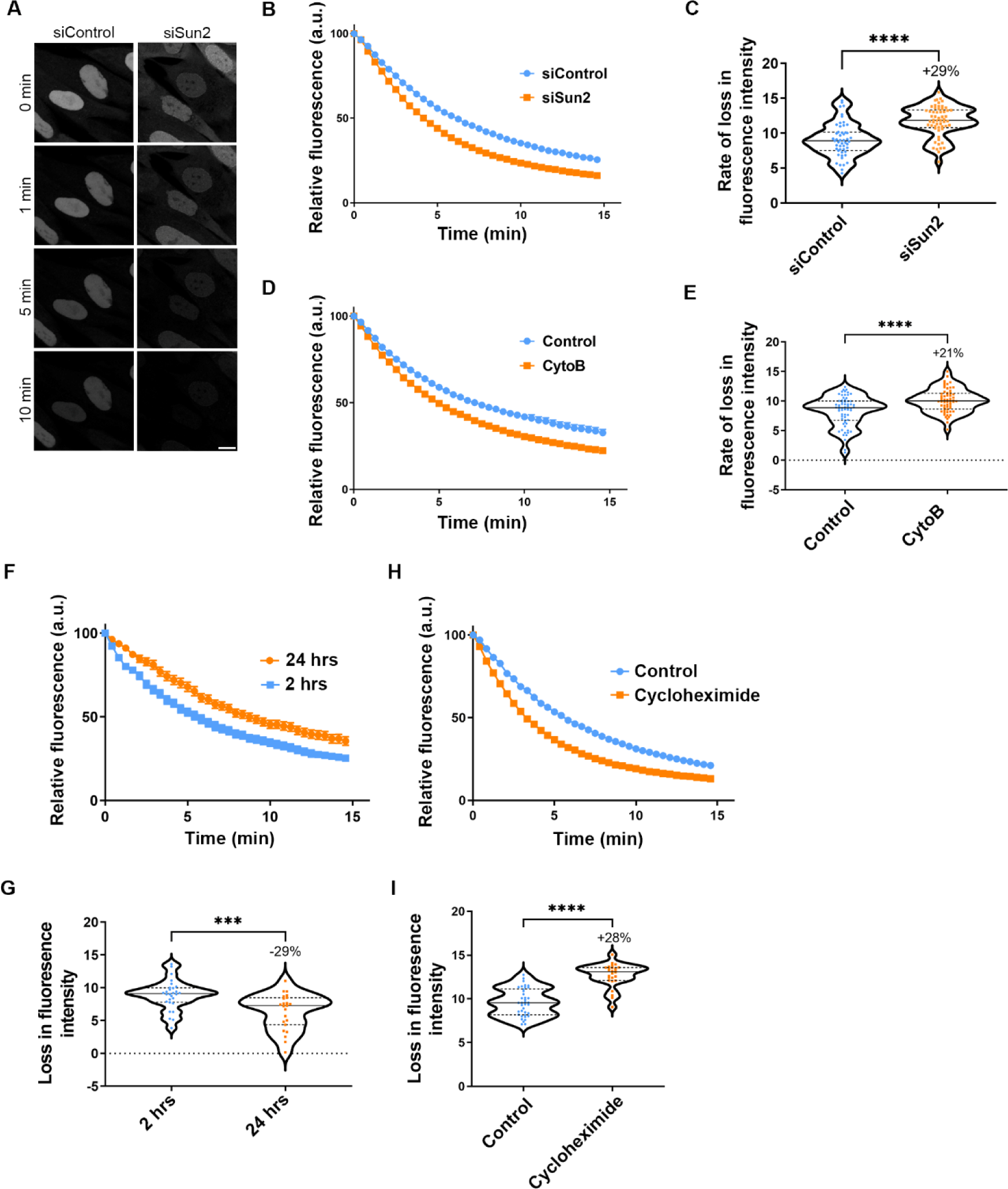
RanGTPase shuttling between the nucleus and cytoplasm is enhanced by conditions that increase available GTP. **(A)** Representative confocal images of GFP-Ran cells depleted of Sun2, where FLIP was used to bleach the cytoplasm and measure the loss of fluorescence intensity from the nucleus. **(B-I)** Graphs of fluorescence change during FLIP **(B, D, F, H)** and rate of loss in fluorescence of GFP-Ran over the first 5 min **(C, E, G, I)** for cells depleted of Sun2 (**(B,C)** siControl n=51, siSun2 n=58; 3 independent replicates), **(D,E)** treated with cytochalasin B (control n=64, cytoB n=60; 3 independent replicates), **(F,G)** plated for 2 or 24 hr (2 hr n=23, 24 hr n=28; 3 independent replicates), or **(H, I)** treated with cycloheximide (control n=38, cycloheximide n=26; 2 independent replicates). Significance calculated using unpaired t-test (C, E, G, I). P***<0.001, P****<0.0001. Scale bar 10 µm. Error bars represent ±SEM.

**Figure 7:**
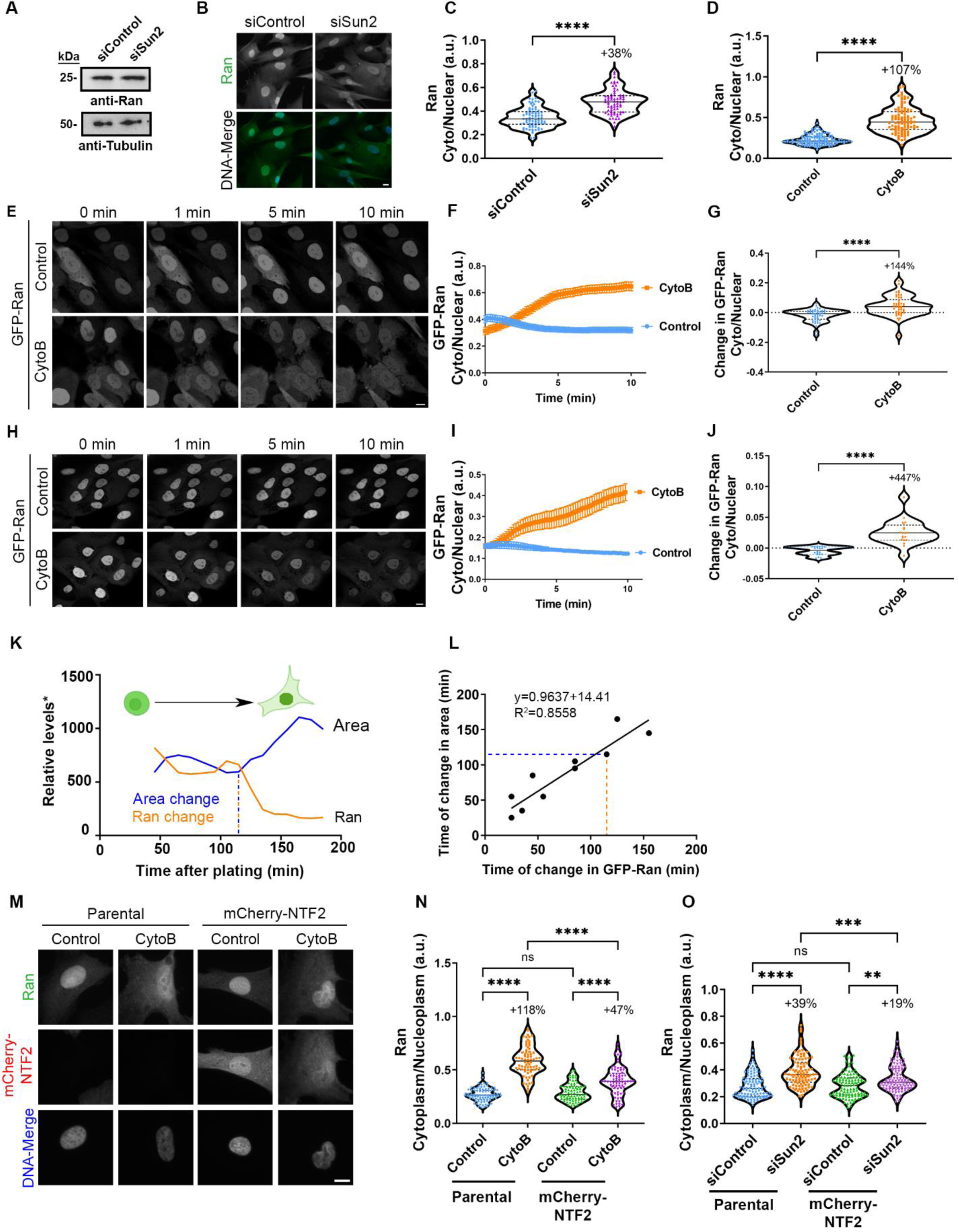
RanGTPase exhibits increased cytoplasmic localization under conditions that elevate levels of available GTP which can be partially reversed by overexpression of NTF2. **(A)** Western blot analysis of BJ-5ta cells depleted of Sun2 and probed for Ran. Tubulin was used as a loading control. **(B)** IF images of Sun2-depleted BJ-5ta cells probed for Ran (green) and stained with Hoechst (blue) to visualize the nucleus. **(C)** Quantification of cytoplasmic to nuclear ratio of endogenous Ran from BJ-5ta cells treated with siSun2 (siControl n=80, siSun2 n=69; 3 independent replicates). **(D)** Quantification of cytoplasmic to nuclear ratio of endogenous Ran in BJ-5ta cells treated with cytochalasin B (control n=130, CytoB n=99; 3 independent replicates). **(E)** Representative confocal images of BJ-5ta cells stably expressing GFP-Ran treated with cytochalasin B at indicated time points. **(F)** Quantification and **(G)** relative rate of change in nuclear to cytoplasmic ratio of GFP-Ran of BJ-5ta cells within the first two min of cytochalasin B treatment (control n= 47, CytoB n=46; 3 independent replicates). **(H)** Representative confocal images of GFP-Ran in MCF10A cells treated with cytochalasin B at indicated time points. **(I)** Quantification and **(J)** relative change in cytoplasmic to nuclear ratio of GFP-Ran in MCF10A cells treated with cytochalasin B (control n=60, cytoB n=60; 3 independent replicates). **(K)** BJ-5ta cells expressing GFP-Ran were trypsinized and plated on fluorodishes for live cell imaging. Representative graph of the relative levels of GFP-Ran and cellular area over time in a single BJ-5ta GFP-Ran cell. The point at which the cellular area or GFP-Ran localization dramatically changes from initial plating is identified as the time of change (indicated with dashed lines).*In order to combine these measurements on a similar scale, cellular area is represented in pixels and Ran localization change is represented by fluorescence (a.u. x 1000). **(L)** Correlation of time of change in GFP-Ran to change in cellular area from (**K**; n=18 across 2 independent experiments). Cells with no change in cellular area were not included in linear regression. **(M)** Representative IF images of BJ-5ta parental or mCherry-NTF2 (red) overexpressing cells treated with cytochalasin B and probed for Ran (green). **(N)** Quantification of endogenous cytoplasmic to nuclear Ran from **(M)** (parental control n=85, parental cytoB n=72, mCherry-NTF2 control n=77, mCherry-NTF2 cytoB n=73; 3 independent replicates). **(O)** Quantification of endogenous cytoplasmic to nuclear Ran in parental or mCherry-NTF2 expressing BJ-5tas depleted of Sun2 (parental siControl n=106, parental siSun2 n=99, mCherry-NTF2 siControl n=104, mCherry-NTF2 siSun2 n=105; 3 independent replicates). Significance calculated using t-test (C, D, G, J), one-way ANOVA with Tukey’s post hoc (N, O). ns P>0.05, P**<0.01, P***<0.001, P****<0.0001. Scale bars 10 µm. Error bars represent ±SEM.

### RNA export and protein synthesis are regulated by altered rates of NCT

Thus far we have demonstrated that naturally occurring or experimentally induced changes in levels of available GTP alter the rates of NCT, but not the outcomes of NCT decisions. We hypothesized that these differences in rates of NCT may have a profound impact on the outcomes of processes that utilize a considerable portion of the cellular transport capacity. RNA export is one such example. Collectively accounting for over 80% of total RNA, rRNA and tRNA utilize Ran-dependent CRM1-mediated nuclear export with the substantially less abundant mRNA predominantly utilizing Ran-independent export pathways (Williams et al., 2018). To assess how altered levels of available GTP modulate the rate of total RNA export, we utilized pulse-chase ethynyl uridine (EU)-based total RNA labeling. After a one-hour labeling, total level of EU-generated signal enables measurement of RNA synthesis. Three hr post-EU washout, the EU signal can be used to assess RNA export from the nucleus by comparing cytoplasmic to nuclear intensity. We first confirmed that blocking CRM1-mediated transport with leptomycin B resulted in an expected shift in RNA localization but not synthesis (Fig. 8, A, B, and C). We next reduced available GTP with MPA and observed a dramatic decrease in RNA export, but not RNA synthesis (Fig. 8, D and E). In contrast, enhancing available levels of GTP by inhibiting protein synthesis resulted in an increase in cytoplasmic RNA but no change in synthesis (Fig. 8 F, G and H). Similarly, increasing GTP levels by LINC perturbation with Sun2 depletion or depolymerization of the actin cytoskeleton with cytochalasin B both led to an increase in RNA export to the cytoplasm, without a change in levels of RNA synthesis (Fig. 8, I, J, K and L). To assess if the steady-state localization of total RNA was altered by enhanced RNA export, we used a live-cell RNA dye and observed a decrease in the ratio of cytosolic to nuclear RNA with inhibition of CRM1-mediated export using leptomycin B or GTP reduction with MPA (Fig. 7, M, N and O). In contrast, we observed an increase in the cytoplasmic levels of RNA following Sun2 depletion, with no change in total RNA levels for any of these conditions (Fig. 7, P, Q and R).

**Figure 8:**
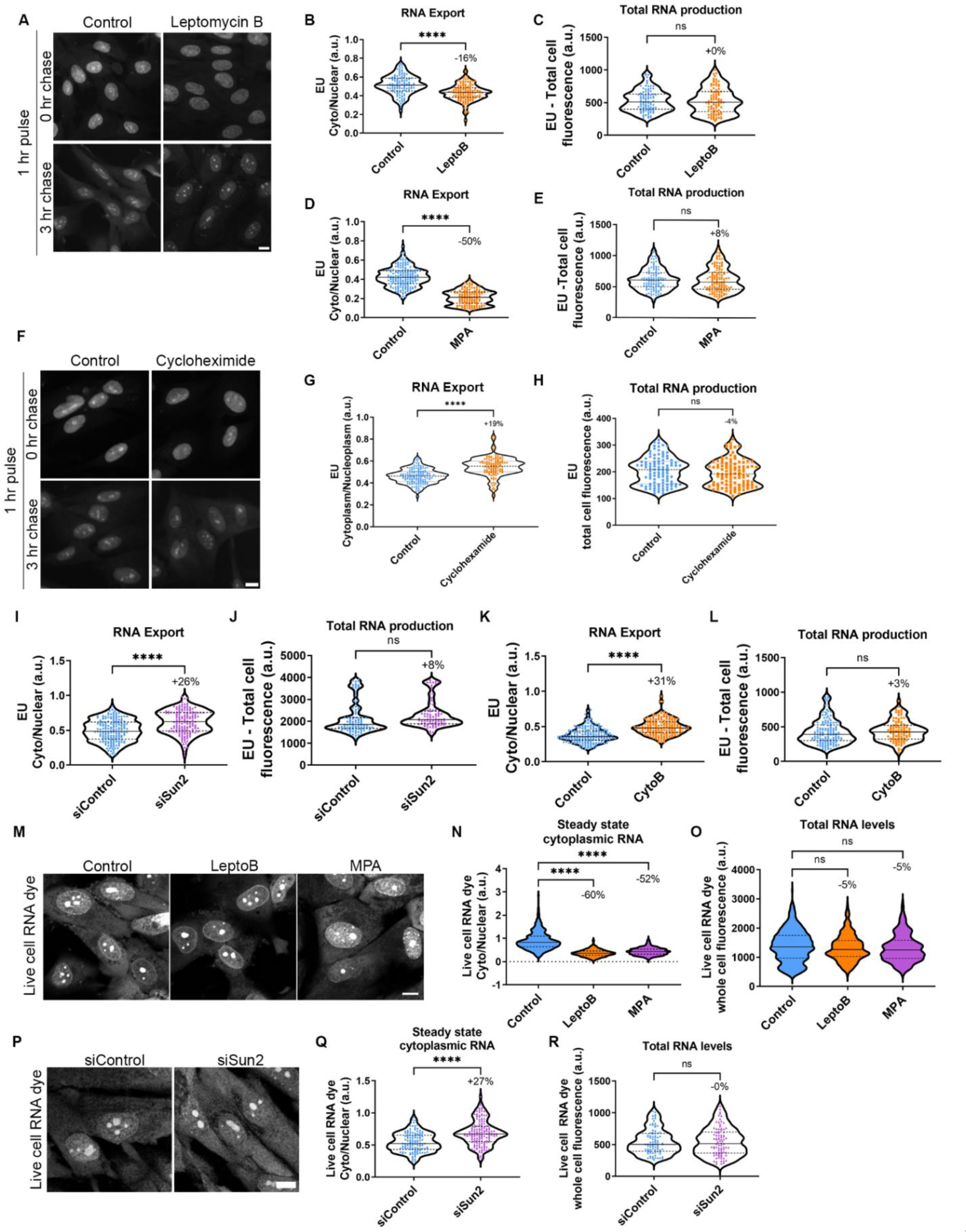
Rates of RNA export and levels of cytosolic RNA are regulated by available GTP. **(A)** Representative IF images of BJ-5ta cells treated with EU for 1 hr (pulse), then re-fed for 0 hr or 3 hr with EU-free media (chase) either untreated (control) or treated with leptomycin B. **(B, C)** Quantification of **(B)** cytoplasmic to nuclear ratio and **(C)** total EU labeling of BJ-5ta cells treated with leptomycin B pulsed with EU for 1 hr, then chased for 3 hr (**B**; control n=97, leptomycin B n=92) or 0 hr (**C**; control n=119, leptomycin B n=128). Results are from 2 independent replicates. **(D, E)** Quantification of **(D)** cytoplasmic to nuclear ratio or **(E)** total EU labeling of BJ-5ta cells treated with MPA pulsed with EU for 1 hr, then chased for 3 hr (**D**; control n=178, MPA n=125) or 0 hr (**E**; control n=111, MPA n=125). Each contain 2 independent replicates. **(F)** Representative IF images of BJ-5ta cells treated with EU for 1 hr (pulse), then refed for 0 hr or 3 hr with EU-free media (chase) treated with cycloheximide. **(G, H)** Quantification of **(G)** cytoplasmic to nuclear ratio or **(H)** total EU labeling of BJ-5ta cells treated with cycloheximide and pulsed with EU for 1 hr, then chased for 3 hr (**G**; control n=146, cycloheximide n=112) or 0 hr (**H;** control n=103, cycloheximide n=106). Each contain 2 independent replicates. **(I, J)** Quantification of **(I)** cytoplasmic to nuclear ratio or **(J)** total EU labeling of BJ-5ta cells depleted of Sun2 pulsed with EU for 1 hr, then chased for 3 hr (**I**; siControl n=175, siSun2 n=161) or 0 hr (**J**; siControl n=92, siSun2 n=79). Each contains 3 independent replicates. **(K, L)** Quantification of **(K)** cytoplasmic to nuclear ratio or **(L)** total EU labeling for cytochalsin B-treated BJ-5ta cells pulsed with EU for 1 hr, then chased for 3 hr (**K**; control n=190, CytoB n=175) or 0 hr (**L**; control n=140, CytoB n=103). Each contains 2 independent replicates. **(M)** Representative confocal images of BJ-5ta cells treated with either MPA or Leptomycin B labeled with live cell RNA dye for 1 hr. **(N,O)** Quantification of **(N)** cytoplasmic to nuclear ratio (control n=582, LeptoB n=517, MPA n=409) or **(O)** total cell fluorescence (control n=606, LeptoB n=536, MPA n=269) across three independent replicates. **(P)** Representative confocal images of BJ-5ta cells depleted of Sun2 labeled with live cell RNA dye for 1 hr. **(Q, R)** Quantification of **(Q)** cytoplasmic to nuclear ratios (siControl n=122, siSun2 n=149) or **(R)** total cell fluorescence (siControl n=86, siSun2 n=86) of live cell RNA dye in cells depleted of Sun2. Each contains 3 independent replicates. Significance calculated using unpaired t-test (B-E, G-J, Q, R) or one-way ANOVA with Dunnett’s post hoc (N and O). ns P>0.05. P****<0.0001. Scale bars 10 µm.

We next investigated if these changes in cytoplasmic RNA levels that result from altered availability of GTP and rates of NCT impact protein synthesis. Utilizing L-Azidohomoalanine (AHA) labeling of newly synthesized proteins to measure relative rates of overall protein translation, we observed an insignificant trend towards reduction of protein synthesis in cells treated with MPA that reduces available GTP and the expected significant decrease with the protein synthesis inhibitor cycloheximide (Fig. 9 A and B). To confirm that a reduction in NCT, and specifically CRM1-mediated export, could significantly reduce protein synthesis, we treated cells with leptomycin B and observed a significant reduction in protein synthesis (Fig. 9, C and D). We next assessed how altering cellular processes that regulate availability of GTP and rates of NCT impact protein synthesis. BJ-5ta cells plated on 1 kPa and 22 kPa substrates had significantly higher levels of protein synthesis compared to cells plated on 46 kPa and 308 kPa substrates, with decreased protein synthesis plateauing at 46 kPa (Fig. 9 E). Finally, we perturbed the LINC complex by Sun2 depletion and observed a significant increase in protein synthesis (Fig. 9 F and G). These studies collectively suggest that altering availability of GTP and rates of NCT positively regulate rates of RNA export, cytoplasmic levels of RNA and the rate of protein synthesis.

**Figure 9:**
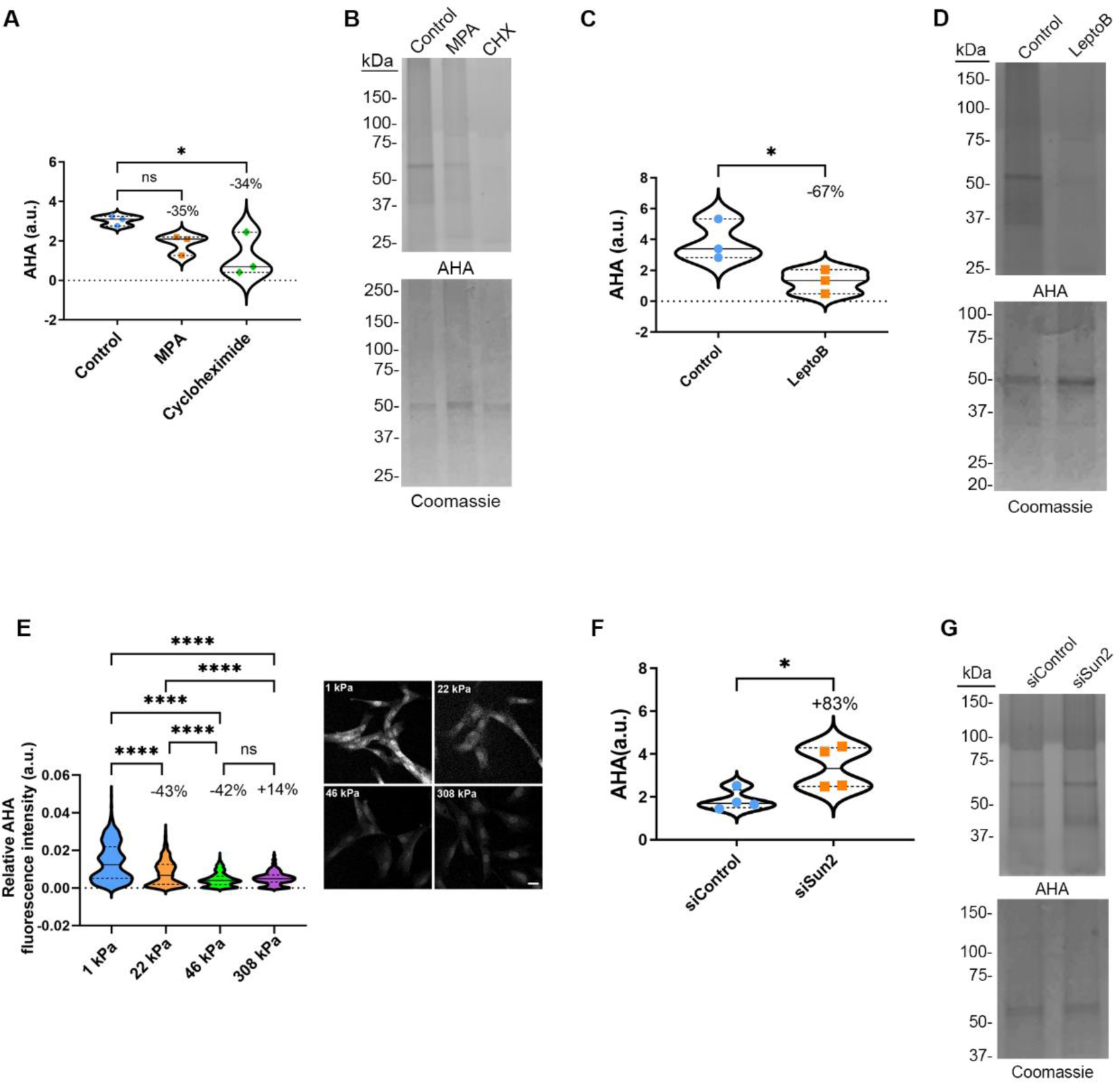
Rates of protein synthesis are regulated by conditions that alter available GTP. **(A, B)** Quantification of protein lysates from AHA-labeled BJ-5ta cells treated with **(A)** MPA or cycloheximide (CHX) and **(B)** analyzed by SDS-PAGE. AHA signal was normalized to Coomassie stain. Results are from three independent replicates. **(C, D)** Quantification of protein lysates from AHA-labeled BJ-5ta cells treated with **(C)** leptomycin B and **(D)** analyzed by SDS-PAGE. AHA signal was normalized to Coomassie stain. Results are from three independent replicates. **(E)** Quantification of protein synthesis and representative images of BJ-5ta cells plated on increasing substrate rigidities, then fixed and labeled with AHA (1 kPa n=646, 22 kPa n=833, 46 kPa n=793, 308 kPa n=530). Results are from three independent replicates. Scale bar 10 µm. **(F,G)** Quantification of protein lysates from AHA-labeled BJ-5ta cells depleted of **(F)** Sun2 and **(G)** analyzed by SDS-PAGE. AHA signal was normalized to Coomassie stain. Results are from four independent replicates. Significance calculated using one-way ANOVA with Dunnett’s post hoc (A), unpaired t-test (C and F) or one-way ANOVA with Tukey’s post hoc (E). ns P>0.05, P*<0.05. P****<0.0001.

## Discussion

Our studies demonstrate that the rate of NCT is regulated by the availability of GTP. Conditions that elevate levels of free GTP, such as inhibition of protein synthesis, reduced cell spreading, functional disruption of actin or microtubule structures, and/or LINC complex perturbation, all lead to an increase in NCT. These energy-dependent effects on NCT appear generalizable to multiple transportin pathways that utilize Ran and impact the dynamic flow of Ran between the nucleus and cytoplasm. Under all conditions examined, altered levels of GTP preserve the need for signal-induced activation of these NCT pathways. This is evident by the need to stimulate glucocorticoid receptor import with dexamethasone even under conditions with enhanced rates of NCT, or the preservation of inducible glucocorticoid receptor import under conditions of reduced NCT. Based on our observations, it is only the rate of NCT that is altered, not the outcome of a signal to activate transport. However, GTP-dependent changes in the rates of NCT can have profound effects on critical biological processes. A reduction in RNA export, induced by conditions that decrease available GTP, also reduces protein synthesis, not necessarily by limiting cytosolic levels of mRNA that largely uses a Ran-independent export pathway (Kohler and Hurt, 2007), but likely by limiting the Ran-dependent transport of rRNA that comprises ∼80% of total RNA (O’Neil et al., 2013) and tRNA that are both needed for ribosome function and thus protein synthesis.

This suggests a possible regulatory feedback loop in which the sensitivity of Ran-mediated transport to GTP levels may function to decrease RNA export and protein translation, a process that substantially consumes cellular GTP (Lindqvist et al., 2018), as a GTP-preserving mechanism within the cell during brief periods of reduced energy levels. However, at this time we cannot rule out that the altered levels of GTP directly impact protein synthesis, although we can demonstrate that direct inhibition of NCT, specifically inhibition of CRM1-mediated export, inhibits protein synthesis without altering available GTP.

We have demonstrated that GTP availability under normal physiological conditions can be a rate-limiting step for NCT, as has been predicted (Gorlich et al., 2003). Impairment of GTP or ATP-consuming processes, such as cytoskeletal-mediated tension generation (DeWane et al., 2021) or protein translation (Lindqvist et al., 2018) lead to enhanced NCT. Since GTP can be generated by NDPK phosphate transfer from ATP (Boissan et al., 2009), GTP levels are intimately related to the availability of free ATP (Schwoebel et al., 2002). Thus, it would be expected that any process that alters cellular energetics would impact NCT. We support the model that it is a combination of Ran’s reduced affinity to GTP as compared to GDP (Klebe et al., 1995b), the similar affinity of RCC1, Ran’s GEF, to RanGTP and RanGDP (Klebe et al., 1995a), and the sheer volume of GTP consumption needed to maintain the Ran gradient, estimated to be in excess of 10^5^ molecules of Ran per second that leave the nucleus (Gorlich et al., 2003; Smith et al., 2002) collectively contribute to the sensitivity of Ran to fluctuations in levels of free GTP. It is unclear if Ran-mediated NCT is uniquely sensitive, or if there are other GTP-dependent processes that are similarly impacted by physiological fluctuations in free GTP levels. It also remains unknown if the transport-independent functions of Ran, such as NE reformation, mitotic spindle assembly, or cytokinesis (Beaudet et al., 2017; Carazo-Salas et al., 2001; Hutchins et al., 2009), are functionally impacted by the relative availability of free GTP in a manner similar to NCT. Our prediction is that cells eventually adapt to an optimal steady-state level of available energy, finding a balance of GTP production sufficient for a sufficient rate of NCT, or perhaps altering mitochondrial abundance or glycolytic pathways if available GTP is insufficient. Transient fluctuations in available GTP, for example due to cell migration or in response to matrix remodeling, would lead to more dramatic and likely acute changes in rates of NCT.

We have found that an elevated rate of NCT correlates with not only increased nucleocytoplasmic shuttling of Ran but also increased levels of cytosolic Ran, presumably due to the inability of NTF2 to reimport RanGDP into the nucleus as rapidly as RanGTP is being removed from the nucleus by recycling importins and exportin-cargo complexes (Beaudet et al., 2017). RanGAP could be a rate-limiting factor for nuclear recycling of Ran since NTF2 binds to RanGDP but not RanGTP. However, limited RanGTP hydrolysis mediated by cytosolic RanGAP, or perhaps simply the binding of RanGTP to the cytosolic RanGAP, would be predicted to reduce NCT since exportin-cargo release and importin-cargo binding both depend on cytosolic conversion of Ran to a GDP bound form and release from the transportin. Additionally, our evidence that NTF2 overexpression at least partially rescues the nuclear enrichment of Ran under normal conditions supports that it is the level of NTF2 that is rate-limiting for the Ran cycle.

Our studies support that rates of NCT are, in general, only indirectly regulated by cellular forces, in as much as generation of those forces decreases free GTP. That is not to say that there are no energy-independent mechanisms that regulate NCT, such as has been proposed via the dilation of NPCs (Zimmerli et al., 2021), but that these do not appear to be the primary mechanism by which cellular forces, including those acting on the nucleus, regulate NCT. Our findings differ from previous studies (Andreu et al., 2022; Elosegui-Artola et al., 2017) that report mechanical forces within the cell enhance rates of NCT due to physical stretching of the NPC, and we are unable to explain this discrepancy as we cannot replicate those findings using some of the same technical approaches. We hypothesize that LINC perturbation leads to elevated levels of available GTP and rates of NCT by mechanisms that could include, directly interfering with the cytoskeletal dynamics and structure (Lombardi et al., 2011; Sharma and Hetzer, 2023), inhibiting nuclear positioning within the cell during migration (Gomes et al., 2005; Luxton et al., 2010; Luxton et al., 2011) and/or by inhibiting the motility of the cell (Lombardi et al., 2011; Sharma and Hetzer, 2023).

The Ran gradient appears exceptionally sensitive to changes in available GTP, which leads not only to altered rates of NCT, but also impacts levels of RNA export from the nucleus and rates of protein synthesis. Bioenergetic regulation of NCT, and thus RNA export and protein synthesis, by titrating the availability of fuel for the Ran gradient is a mechanism by which cells can convert energy fluctuations, including those generated by cellular processes such as migration or spreading, into physiological changes that could impact key processes such as cell cycle, differentiation, and metabolism. It is also tempting to speculate that perhaps some of the defects in NCT that have been reported for various neurodegenerative diseases could be related to altered bioenergetics (Strope et al., 2022), and/or that the altered bioenergetics that are proposed to underlie several neurodegenerative diseases (Ding and Sepehrimanesh, 2021) could lead to defective NCT as a contributory mechanism of disease. Perhaps post-mitotic cells with a reduced capacity for adaptation would be more sensitive to a bioenergetic decline and be susceptible to reduced NCT.

## Author Contributions

Conceptualization, K.L.S, T.P.L, & K.J.R; Methodology, K.L.S, C.T.H, A.D.H, & K.J.R; Investigation, K.L.S, C.T.H, A.D.H; Resources, P.P, T.P.L; Writing – Original Draft, K.L.S, T.P.L, & K.J.R.; Writing – Review & Editing, K.L.S, C.T.H, A.D.H, T.P.L, & K.J.R.; Visualization, K.L.S, C.T.H, A.D.H, & K.J.R; Funding Acquisition, T.P.L & K.J.R; Supervision, K.J.R

## Methods

### Plasmid production

All plasmids were constructed using the Takara Infusion cloning kit. NES-mCherry-LINuS pBabe puro and pBabe neo were constructed using pDN77 (Gift from Barbara Di Ventura & Roland Eils; Addgene plasmid # 61347) as a template amplified with PCR using primers (5’)TGGTACGTAGGAATTCGCCACCATGCCCAGCACCCGG and (5’) ATTCCACAGGGTCGACCTAGTCCAGCTTTTTCTTCTTGGCTG. This was recombined with mCherry-NLS pBabe neo (Halfmann et al., 2019) digested with EcoRI and SalI. GFP-KASH4 was constructed as previously described (Roux et al., 2009). GFP-Ran pBabe puro was constructed using human Ran WT cDNA (gift from Brian Burke) as a template amplified with primers (5’) CTGTACAAGGACCTCGAGGCTGCGCAGGGA and (5’) ACATTCCACAGGGTCGACTCACAGGTCATCATCC. This was recombined with GFP-BAF pBabe puro (Halfmann et al., 2019) digested with XhoI and SalI. mCherry-NTF2 pBabe neo was constructed using pET23b-NTF2 (gift from Larry Gerace; Addgene plasmid # 108918) as a template amplified with primers (5’) TCCGGACTCAGATCTCGAGCAATGGGAGACAAGCCAA and (5’) ATTCCACAGGGTCGACTCAGCCAAAGTTGTGC. This was recombined with mCherry-NLS pBabe neo digested with XhoI and SalI. GR-GFP pBabe puro was constructed using pk7-GR-GFP (Gift from Ian Macara; Addgene plasmid # 15534) amplified with primers (5’) AGTGTGGTGGTACGTAGCCACCATGGACTCCAAAGAAT and (5’) CCGCTCGACGACAGGGGCCCCTTTTGATGAAACAGA. This was recombined with Lap2β-GFP pBabe puro (Halfmann et al., 2019) digested with SnaBI and ApaI. GFP pBabe puro was constructed by amplifying GFP with primers (5’) TGGTACGTAGGAATTCTAGCCACCATGGTGAGCA and (5’) TTACTTGTACAGCTCGTCCATGCCCTCGAGACATTCCACAGGGTCGAC and recombined with pBabe puro digested with EcoRI and XhoI. GEVALNull pLVp_puro and GEVAL30 pLVp_puro were gifts from Mikhail A. Nikiforov. KRAB-dCas9-IRES-NES-LINuS-mCherry pBabe neo used for CRISPRi-mediated knockdown was constructed using pLX_311-KRAB-dCas9 (a gift from John Doench & William Hahn & David Root; Addgene plasmid # 96918) amplified using primers (5’) AGTGTGGTGGTACGTAGCCACCATGGATGCTAAGTCAC and (5’) CGGAATTGATCCCGCGAATTCTTAGGATCCGCTGCTG. This was recombined into SnaBI and EcoRI digested IRES-NES-LINuS-mCherry pBabe neo. The IRES-NES-LINuS-mCherry pBabe neo backbone vector was created by amplifying an internal ribosome entry site (IRES) from pMSCV-IRES-mCherry-FP (a gift from Dario Vignali, Addgene plasmid #52114) using primers (5’) AGCATCATGTAAGAATTCGCGGGATCAATTCCG and (5’) CTTGCTCACCATGTTTAAACTTTATCGTGTT and recombining into EcoRI digested NES-mCherry-LINuS pBabe neo. The plasmids ss-GFP-L-KDEL and ss-Sun1-L-KDEL pcDNA3.1 were constructed as previously described (Zhang et al., 2019). GFP-NLSx3 pBabe puro was constructed as previously described (Halfmann et al., 2019).

### Cell culture

BJ-5ta cells were obtained from American Type Cell Culture (ATCC; CRL-4001) and cultured in Dulbecco’s Modified Eagle Medium (DMEM) with 4.5 g/L glucose, sodium pyruvate, and L-glutamine (Corning) supplemented with 10% Fetal Bovine Serum (FBS). MCF10A cells were obtained from ATCC (CRL-10317) and cultured in Mammary Epithelial Cell Growth Media (Lonza) supplemented with MEGM Mammary Epithelial Cell Growth Medium BulletKit (Lonza), 10% FBS, and cholera toxin (100 ng/mL). Gentamicin sulfate-amphotericin was omitted from the media. Cells were maintained at 37°C and 5% CO2. pBabe stable cell lines were generated using retroviral transduction. HEK293 Phoenix cells were transfected with the plasmid using Lipofectamine 3000 (ThermoFisher Scientific). The transfected cells were incubated at 37°C for 5 hr. The media was changed and the cells were incubated at 32°C for 48 hr. The media was then passed through a 0.45 µM filter onto target cells with polybrene (2.5 µg/mL). After 48 hr, either puromycin (ThermoFisher Scientific, 0.5 µg/mL) or G418 sulfate (Corning, 50 µg/mL) was added to cells to select for the plasmid of interest. Puromycin was used for pBabe puro plasmids for 2-3 days and G418 sulfate was used for pBabe neo plasmids for 5-7 days. GEVAL stable cell lines were generated by the protocol above using third-generation lentiviral plasmids (CellBio labs) transfected in 293-T cells (ATCC CRL-3216). All cells were tested monthly for mycoplasma contamination. GFP pcDNA3.1, GFP-KASH4 pcDNA3.1, ss-GFP-KDEL pcDNA3.1, and ss-Sun1L-GFP-KDEL pcDNA3.1 were transiently transfected into cells stably expressing LINuS using Lipofectamine 3000 (ThermoFisher Scientific) when cells were 80% confluent. Media was changed 6 hr after transfection. LINuS studies were performed 48 hr after transfection. CRISPRi-mediated depletion of various LINC-complex constituents was performed using a KRAB-dCas9 system (Rosenbluh et al., 2017).

### Drug Treatments

Cells were treated with cytochalasin B (Tocris Biosciences, 20 µM, 10 min) and latrunculin B (MilliporeSigma, 1 µM, 10 min) to depolymerize actin, and nocodazole (MilliporeSigma, 1 µg/mL, 15 min) to depolymerize microtubules. Cells were treated with leptomycin B (Sigma-Aldrich, 20 ng/mL, 48 hr) to inhibit CRM1-mediated export. Importazole (Selleck Chemicals, 40 µM, 1 hr) was used to inhibit Importin β activity. Cells were treated with guanosine (TCI America, 100 µM, 48 hr) to supplement GTP synthesis. Cells were treated with cycloheximide (ThermoScientific, 350 µM, 2 hr) to inhibit protein translation. Cells were treated with MPA (ThermoScientific, MPA, 1.8 µM, 48 hr) to deplete GTP. Cells were treated with sodium azide (Sigma-Aldrich, 10mM, 20 min) and 2-deoxyglucose (Sigma-Aldrich, 6mM, 20 min) to deplete ATP. Control cells were treated with equal volumes of media. Nuclear import of GR-GFP was induced by the addition of dexamethasone (Sigma-Aldrich, 1 µM, 15 min).

### Scratch Assay

MCF10A cells expressing LINuS, GEVALNull, or GEVAL30 were plated on fluorodishes and allowed to attach for 48 hr. A 200 µL pipette tip was then scraped along the bottom of the fluorodish to create a cross-shaped ‘scratch wound’ in the cell monolayer, followed by a PBS rinse and re-feed with fresh media to remove denuded cells. The cells were allowed to migrate into the denuded area for 2 hr prior to imaging.

### RNA Studies

Ethynyl Uridine (Invitrogen, 1 mM) was incubated with cells in media for one hr, prior to fixation (0 hr chase), or the EU containing media was replaced and cells were incubated for 3 hr, prior to fixation (3 hr chase). Cells were fixed with 3% w/v PFA in PBS for 10 min then permeabilized with 0.4% w/v Triton-X 100 in PBS for 15 min. EU was visualized using the Click-It cell reaction buffer kit (Invitrogen) and AlexaFluor 488 Azide (Invitrogen) according to manufacturer instructions. Cells at 80% confluency were incubated with live-cell RNA dye (Live Cell RNA Imaging Kit, 400X stock; Cell Navigator) for 1 hour in media. Images were captured as described below and quantified using ImageJ.

### Translation assay

Cells were grown in methionine-free media (Gibco) for one hour, then L-Azidohomoalanine (AHA) was added to the cells for 6 hr. For SDS-PAGE analysis, cells were collected, lysed, sonicated, and the Click-It Protein reaction (Invitrogen) was performed with Tetramethylrhodamine (TAMRA) alkyne (Invitrogen) according to manufacturer’s suggestions; however, the methanol precipitation was omitted and samples were analyzed by SDS-PAGE immediately following Click-It labeling. Gel images were acquired on a Licor Odyssey system for AHA fluorescence signal. Subsequently, the gels were stained with Coomassie Brilliant Blue R250 and imaged on a scanner. AHA fluorescence was normalized to Coomassie stain for each sample. For *in cellulo* AHA labeling, cells were fixed with 3% w/v PFA in PBS for 10 min then permeabilized with 0.4% w/v Triton-X 100 in PBS for 15 min. AHA was visualized using the Click-It cell reaction buffer kit (Invitrogen) and TAMRA alkyne (Invitrogen) according to manufacturer instructions. Fixed and labeled cells were imaged on a Nikon A1R confocal microscope (see below for details). Quantification of AHA labeling in fixed cells on substrates was performed with Cell Profiler v 4.2.4, using a pipeline that measured average AHA intensity with Otsu thresholding and a correction factor between 0.5-1.0.

### siRNA and CRISPRi experiments

siRNA and sgRNA transfections were performed using RNAiMax (ThermoFisher) according to the manufacturer’s instructions. Cells were seeded at 70% confluency into 12-well tissue-culture plates containing 1 mL DMEM and allowed to adhere to the bottom of the plate during an overnight incubation at 37 °C. Cells were transfected 24 and 72 hr after being seeded, then trypsinized 96 hr after being seeded and replated on FluoroDishes for 48 hr prior to imaging. ON-TARGETplus SMARTpool siRNAs (Horizon Discovery) were used to target Sun1 (NM_001130965.3), Sun2 (NM_001199579.2), vimentin (NM_003380.5), nesprin1 (NM_001347701.2), nesprin2 (NM_015180.6), and nesprin3 (NM_152592.6). 15 pmol siRNA (0.75 µl of a 20 µM solution in RNase free H_2_O) were used per transfection in all knockdowns. For Sun1/2 co-depletion, total siRNA oligo concentration were as described above. CRISPRi guide RNAs (sgRNAs; Horizon Discovery) were used to target Sun1 (CI-025277-01-0002), Sun2 (CI-009959-01-0002), vimentin (CI-003551-01-0002), Nesprin1 (CI-014039-01-0002), and Nesprin2 (CI-019259-01-0002). CRISPRi-mediated knockdowns were performed using 25 pmol sgRNA (1.25 µl of a 20 µM solution in RNase free H_2_O) per transfection. For Sun1/2 co-depletion, total sgRNA oligo concentration were as described above. All knockdowns for siRNA and CRISPRi were done in parallel with non-targeting control oligos. All siRNA transfections were validated with Western Blot analysis or immunofluorescence.

### Polyacrylamide substrate synthesis and functionalization

Polyacrylamide gels of four different stiffness were synthesized using an established protocol in literature (Munevar et al., 2001). For obtaining desired stiffness of 1, 22, 46 and 308 kPa, acrylamide (Bio-Rad) and bis-acrylamide (Bio-Rad) were mixed in 50:1, 40:1, 20:1 and 12.5:1 ratios (v/v) respectively, to obtain polymer solutions. Polyacrylamide gel solutions were prepared by adding 99.4% v/v of polymer solution (of corresponding stiffness) with 0.5% v/v ammonium per-sulfate (ThermoFisher Scientific) and 0.1% v/v TEMED (ThermoFisher Scientific). Each glass-bottomed fluorodish was treated with bind silane for 5 min to make the bottom hydrophilic. 35 µl polyacrylamide gel solution was sandwiched between the hydrophilic fluorodish and a hydrophobic glass coverslip (18 mm diameter) for ∼20 min at room temperature to complete the polymerization. After the gel polymerized into a uniform layer, the hydrophobic coverslip was peeled off and the gel was hydrated in PBS for 30 min. Following hydration, the gels coated with Sulfo-SANPAH (G-Biosciences) were functionalized using high intensity UV for 216 s. The residual Sulfo-SANPAH was removed by washing the gels with PBS three times at room temperature and the gels were further sterilized under low intensity UV for 30 min. The sterilized gels were coated with fibronectin (10 µg/ml) overnight at 4°C before seeding cells.

### Immunofluorescence

Cells were grown on glass coverslips and fixed with 3% w/v PFA in PBS for 10 min then permeabilized with 0.4% w/v Triton-X 100 in PBS for 15 min. Cells were incubated with primary antibodies for 30 min: Mouse anti-Ran (BD Transduction Laboratories, 610341, 1:100), rabbit anti-Nesprin1 (Atlas Antibodies, HPA019113, 1:100), rabbit anti-Nesprin2 (Atlas Antibodies, HPA003435, 1:50), rabbit anti-Nesprin2/4 (YenZyme; peptide antigen KKAELEWDPAGDIGGLGPLGQ, 1:25, (Neelam et al., 2015)), and rabbit anti-Nesprin3 (Atlas Antibodies, HPA07740, 1:50). Primary antibodies were detected with anti-mouse AlexaFluor 488 (ThermoFisher Scientific, A11029, 1:1000), anti-Rabbit AlexaFluor 488 (ThermoFisher Scientific, A11034, 1:1000), and DNA was visualized with Hoescht 33342 (1:5000). Coverslips were mounted using 10% (w/vol) Mowiol 4-88 (Polysciences). Fluorescence images were captured using a Nikon Eclipse NiE (40×/0.75 Plan Fluor Nikon objective; 20×/0.75 Plan Apo Nikon objective) microscope at room temperature with a charge-coupled device camera (CoolSnap HQ; Photometrics, Tucson, AZ, USA) linked to a workstation running NIS-Elements software (Nikon, Melville, NY, USA). All images were processed in Adobe Photoshop CC 2017 (Adobe, San Jose, CA, USA) for cropping and brightness/contrast adjustment when applicable.

### Western Blot

Cells were lysed in SDS-PAGE sample buffer, boiled for 5 min, and then sonicated to shear DNA. Samples were separated on 4-20% gradient gels (Mini-PROTEAN TGX; BioRad), then transferred to a nitrocellulose membrane. Membranes were blocked using 10% (vol/vol) adult bovine serum and 0.2% TritonX-100 in PBS for 20 min. The membrane was incubated with primary antibodies (anti-Sun1 (Atlas Antibodies, HPA008346, 1:1000), rabbit anti-Sun2 (Atlas Antibodies, HPA001209, 1:1000), mouse anti-Vimentin (Abcam, ab8978, 1:1000), mouse anti-Ran (BD Transduction Laboratories, 610341, 1:5000), and mouse anti-tubulin (SantaCruz Biotechnology, DM1A, 1:10,000). Primary antibodies were labeled using goat anti-rabbit HRP (ThermoFisher Scientific, G21234, 1:10,000) and goat anti-mouse HRP (Thermofisher Scientific, A24524, 1:10,000), and detected using enhanced chemiluminescence via a Bio-Rad ChemiDoc MP System (Bio-Rad, Hercules, CA, USA). Between each primary antibody, the membrane was quenched using 30% H_2_0_2_.

### Live Cell Imaging

LINuS, LEXY, GFP-Ran, GEVAL, and GR-GFP imaging were performed on a Nikon A1R confocal microscope (Plan Apo λ 20x/0.75, Plan Fluor 40x/1.30 Oil DIC H N2, Apo 60x/1.30 Oil λS DIC N2) with a charge-coupled device camera (CoolSnap HQ; Photometrics, Tucson, AZ, USA) linked to a workstation running NIS-Elements software (Nikon, Melville, NY, USA) with a heated and CO_2_ incubated chamber. Cells were seeded onto 35-mm glass-bottom FluoroDishes (World Precision Instruments) or 8 well glass bottomed chamber slides (Thermofisher Scientific) and were imaged in phenol red-free DMEM supplemented with 10% FBS. Cells were equilibrated in the live cell chamber for 15 min prior to imaging. All images were processed in Adobe Photoshop CC 2017 (Adobe) for cropping and brightness/contrast adjustment when applicable. LINuS cells were allowed to equilibrate in the dark for 10 min prior to activation. LINuS was photoactivated with a 488 nm laser at 100% laser power for 1 second every 30 seconds for 3.5 min. An image was captured immediately after each photoactivation. For the recovery stage, images were captured every 30 seconds for 3 min. Cells that exhibited no import or export of LINuS were excluded from analysis. Nuclear and cytoplasmic LINuS intensity was measured using ImageJ. The relative nuclear signal was calculated using the ratios of nuclear to cytoplasmic intensity of LINuS cells, and starting ratios (just prior to photoactivation) were adjusted to 1. Import and export rates were calculated by finding the slope of the line for the activation and recovery phases, respectively, as described in Figure 1C. GEVAL images were acquired using dual excitation/emission imaging and capturing images at Ex405/Em530 and Ex488/Em530, as previously described in Bianchi-Smiraglia and Rana, 2018. ImageJ was used to calculate the mean fluorescence of nuclei within individual cells for each channel. Ratiometric images were generated using the Image calculator function on ImageJ.

Fluorescence loss in photobleaching (FLIP) experiments were performed by repeatedly bleaching the cytoplasm of GFP-Ran expressing cells for 1 second immediately followed by image capture, every 16 seconds for 15 min with a 488 nm laser at 100% laser power. Fluorescence intensity of the nucleus and cytoplasm was measured with ImageJ. A non-bleached cell in the same frame was used as a control for photobleaching. Significance was calculated by comparing the rates of fluorescence loss in the first five min of photobleaching.

### Nuclear shape analysis

Nuclear volume and height measurements were calculated using consecutive confocal Z-stack images every 0.3 µm of nuclei stained with DAPI using the 60X objective. Nuclear volume measurements were calculated by taking the sum of the area of each stack. The nuclear height was calculated by multiplying the total number of stacks by the distance between each stack.

### Statistical analysis

Unpaired t-test, one-way ANOVA with Dunnett’s post hoc multiple comparison test, or one-way ANOVA with Tukey’s post hoc multiple comparison test were used to find significance, as denoted in the figure legends. Outliers were identified prior to analysis using ROUT analysis with Q set to 1%. Graphs with error bars exhibit mean values ± SEM. For violin plots, solid lines represent the median and dashed lines represent quartiles. Percent change represents comparative change in mean. Statistical analysis, outlier identification, and import and export rates graph generation were performed in GraphPad Prism v.9.5.1 (GraphPad Software Inc., San Diego, CA, USA).

## Acknowledgements

This work was supported by R35GM126949 to K.J.R. from the National Institutes of Health. The Sanford Research Histopathology and Imaging Core and Flow Cytometry Core, which facilitated these studies, is supported by Institutional Development Awards from the National Institute of General Medical Sciences and the National Institutes of Health under grant P20GM103548. Tanmay P. Lele, CPRIT Scholar in Cancer Research, acknowledges support from NIH U01 CA225566 and CPRIT Established Investigator Award RR200043.

The authors declare no competing financial interests.

