## Supplemental Figures for "Nucleocytoplasmic transport rates are regulated by cellular processes that modulate GTP availability"

### Supplementary Figures

**A**

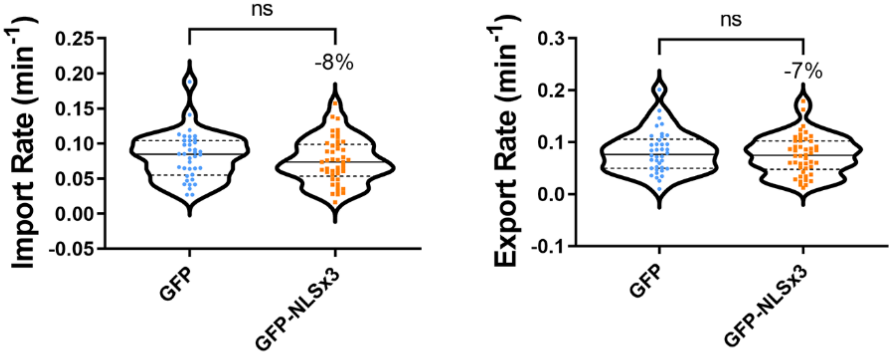


Supplementary Figure 1: Assessing the impact of overexpression of an NLS-containing protein on NCT. **(A)** Import and export rates of BJ-5ta cells expressing LINuS and GFP (n=38) or GFP-NLSx3 (n=46). Results are from two independent replicates. Significance calculated using unpaired t-test. ns p>0.05.

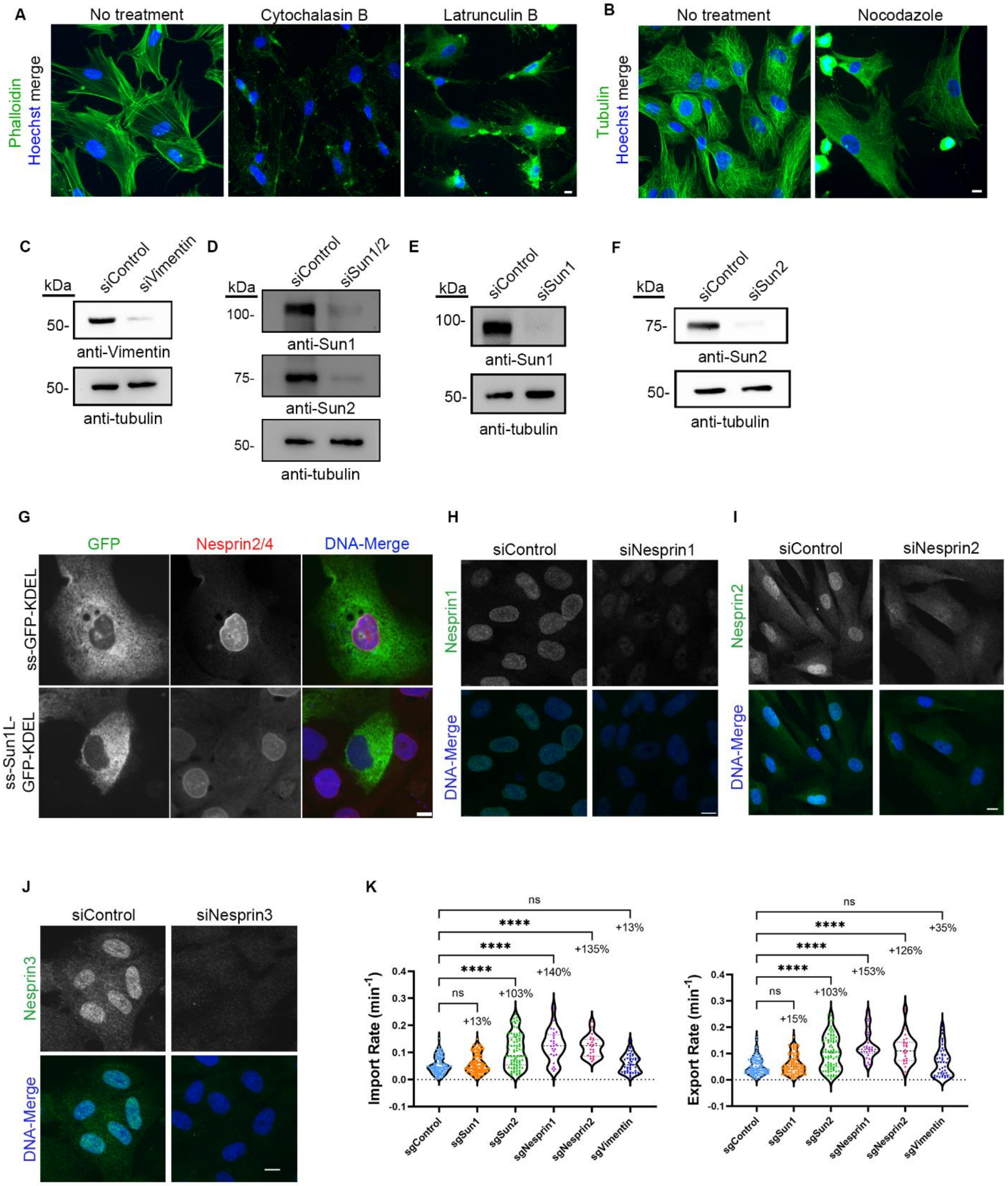

A
No treatment
Cytochalasin B
Latrunculin B
Phalloidin Hoechst merge
B
No treatment
Nocodazole
Tubulin Hoechst merge
C
siControl
siVimentin
kDa
50-
anti-Vimentin
50-
anti-tubulin
D
siControl
siSun1/2
kDa
100-
anti-Sun1
75-
anti-Sun2
50-
anti-tubulin
E
siControl
siSun1
kDa
100-
anti-Sun1
50-
anti-tubulin
F
siControl
siSun2
kDa
75-
anti-Sun2
50-
anti-tubulin
G
GFP
Nesprin2/4
DNA-Merge
ss-GFP-KDEL
ss-Sun1L-GFP-KDEL
H
siControl
siNesprin1
Nesprin1
DNA-Merge
I
siControl
siNesprin2
Nesprin2
DNA-Merge
J
siControl
siNesprin3
Nesprin3
DNA-Merge
K
Import Rate (min$^{-1}$)
Export Rate (min$^{-1}$)
ns
****
ns
+13%
+103%
+140%
+135%
+15%
+153%
+126%
+35%
sgControl
sgSun1
sgSun2
sgNesprin1
sgNesprin2
sgVimentin
0.4
0.3
0.2
0.1
0.0
-0.1

Supplementary Figure 2: Validation of drug treatments and LINC complex disruption. **(A)** IF validation of actin-cytoskeleton disruption in BJ-5ta cells treated with either cytochalasin B or latrunculin. Phalloidin (green) was used as a marker for F-actin. **(B)** IF validation of tubulin disruption in BJ-5ta cells treated with nocodazole. **(C-F)** Western blot validation of siRNA knockdowns of **(C)** vimentin **(D)** Sun1 and Sun2 (Sun1/2), **(E)** Sun1, or **(F)** Sun2 knockdown in BJ-5ta cells. Tubulin was used as a loading control. **(G)** Representative fluorescence images of MCF10A cells transiently transfected with ss-GFP-KDEL or ss-Sun1L-GFP-KDEL (green) probed for Nesprin2/4 (red) to assess loss of LINC complex nesprins from the NE. **(H-J)** Representative fluorescence images of BJ-5ta cells depleted of **(H)** Nesprin1, **(I)** Nesprin2, or **(J)** Nesprin3 (green), labeled with respective antibodies. Hoescht was used to visualize DNA (blue). Scale bars 10 µm. **(K)** Import and export rates of BJ-5ta cells expressing KRAB-dCas9-IRES-LINuS (CRISPRi) and transfected with gRNAs against Sun1 (sgSun1), Sun2 (sgSun2), Nesprin1 (sgNesprin1), Nesprin2 (sgNesprin2), and vimentin (sgVimentin). sgControl (n=109), sgSun1 (n=106), sgSun2 (n=80); 3 independent replicates. sgVimentin (n=42), sgNesprin1 (n=25), sgNesprin2 (n=24); 2 independent replicates. Significance calculated using one-way ANOVA with Dunnett's post hoc. ns P>0.05, P***<0.001, P****<0.0001.

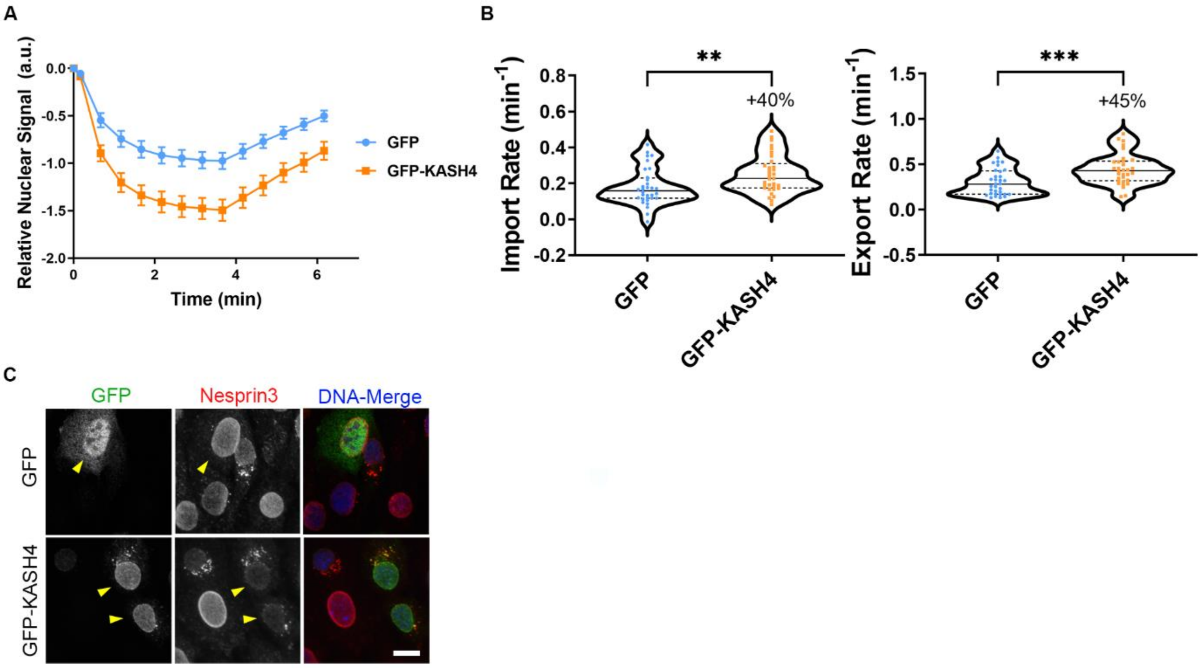


Supplementary Figure 3: Validation of LINC perturbation on NCT using the LEXY reporter. **(A)** Quantification of LEXY nuclear localization during photoactivation in MEFs transiently transfected with either GFP or GFP-KASH4 to disrupt the LINC complex (GFP n=32, GFP-KASH4 n=33). Results are from three independent replicates. **(B)** Import and export rates from A. **(C)** Representative fluorescence images of MEFs transfected with either GFP or GFP-KASH4 (yellow arrows) labeled with antibodies against Nesprin3 to validate nesprin mislocalization from the NE in the presence of dominant-negative GFP-KASH4. Scale bar 10 µm.
